# Cell cycle dynamics of lamina associated DNA

**DOI:** 10.1101/2019.12.19.881979

**Authors:** Tom van Schaik, Mabel Vos, Daan Peric-Hupkes, Bas van Steensel

**Affiliations:** Division of Gene Regulation and Oncode Institute, Netherlands Cancer institute, Amsterdam, the Netherlands

## Abstract

In mammalian interphase nuclei more than one thousand large genomic regions are positioned at the nuclear lamina (NL). These lamina associated domains (LADs) are involved in gene regulation and may provide a backbone for the overall folding of interphase chromosomes. While LADs have been characterized in great detail, little is known about their dynamics during interphase, in particular at the onset of G1 phase and during DNA replication. To study these dynamics, we developed an antibody-based variant of the DamID technology (named pA-DamID) that allows us to map and visualize genome – NL interactions with high temporal resolution. Application of pA-DamID combined with synchronization and cell sorting experiments reveals that LAD – NL contacts are generally rapidly established early in G1 phase. However, LADs on the distal ∼25 Mb of most chromosomes tend to contact the NL first and then gradually detach, while centromere-proximal LADs accumulate gradually at the NL. Furthermore, our data indicate that S-phase chromatin shows transiently increased lamin interactions. These findings highlight a dynamic choreography of LAD – NL contacts during interphase progression, and illustrate the usefulness of pA-DamID to study the dynamics of genome compartmentalization.

## Introduction

The nuclear lamina (NL) is a protein layer underneath the inner nuclear membrane, consisting of lamins and a variety of other proteins. The NL is thought to serve as an anchoring platform for the genome. DNA contacts the NL through large genomic regions named lamina-associated domains (LADs) [1]. Mammalian cells have approximately one thousand LADs that are distributed across the genome. LADs have a median size of ∼0.5 Mb and jointly cover 30-50% of the genome. LADs are associated with gene repression and have been hypothesized to form a backbone for genome organization [2–7].

When dividing cells enter prophase, LADs lose their NL contacts concomitant with chromosome condensation [8]. In metaphase, the NL is disassembled and most of its proteins are excluded from the chromatin [9, 10]. After completion of mitosis, the NL reforms around the still-condensed DNA. B-type lamins localize at the surface of the decondensing chromosomal mass, while Lamin A is initially present throughout the chromatin and only later concentrates at the nuclear periphery [11, 12].

How LAD – NL contacts are re-established during telophase and early interphase is poorly understood. From a genomic perspective, it is not known whether all LADs engage equally in these early contacts, or whether a specific subset of LADs acts as early nucleation sites, with the remainder following later. Microscopy studies found that telomeres are enriched near the NL in early G1 phase, leading to the hypothesis that telomeres may assist in NL reassembly onto chromatin [13]. Whether other chromosomal regions also engage in early contacts with the NL is not known, because systematic analysis of these contacts has not been possible. Similarly, it is not known how the NL contacts of individual LADs are established. For example, it is conceivable that these contacts initiate at specific sequences within LADs, such as LAD boundaries or internal sites, and then gradually spread to cover entire LADs. Testing such models requires mapping of LAD - NL contacts with high temporal resolution.

It is also poorly understood how LAD - NL contacts develop throughout interphase. Microscopy studies have shown that chromatin is relatively mobile early after cell division, but a few hours into interphase this mobility becomes substantially constrained [14–16] and LADs show only movements over a range of ∼0.5 µm [8, 17]. It may thus be expected that LAD - NL contacts do not change much after their establishment, but this has not been investigated by genome-wide analysis.

Similarly, little data is available on the dynamics of LADs during S-phase. This is of interest, because several links have been reported between the NL and DNA replication. Mid-to-late replicating DNA is concentrated near the NL, and genome-wide maps show that late-replicating DNA overlaps strongly with LADs [18, 19]. Furthermore, during S-phase B-type lamins have been found to transiently overlap with replication foci in the nuclear interior, at least in some cell types [20]. Other studies have indicated that lamins are important for DNA replication [reviewed in 21]. These observations raise the interesting question whether interactions of LADs with the NL are subject to changes during S-phase.

So far, the cell cycle dynamics of genome - NL interactions have primarily been studied by microscopy. While these studies have been highly informative, they were often limited to a few selected loci. Hence, a genome-wide view of the dynamics of LAD - NL interactions is still lacking. For genome-wide mapping of NL interactions, DamID has been the major method [1, 22]. DamID is based on expression of a fusion protein of Dam and a NL protein (e.g. Lamin B1), which results in gradual accumulation of adenine methylation (^m6^A) on DNA that contacts the NL [23]. This ^m6^A labeled DNA is then amplified and sequenced. An added advantage of DamID is that the ^m6^A tags can be detected by a GFP-labeled ^m6^A-Tracer protein [8]. This enables visualization of LAD – NL contacts *in situ*, which greatly assists in the interpretation of genome-wide DamID mapping data.

However, a major limitation of DamID is its poor temporal resolution. The activation of Dam and deposition of ^m6^A requires at least several hours [8, 24], precluding detailed analysis of the NL interaction dynamics. To overcome these limitations, we developed pA-DamID – a hybrid of DamID and the CUT&RUN method [25]. This allows for both mapping and visualization of NL contacts with high temporal resolution. Using pA-DamID, we show that after mitosis NL contacts do not initiate at defined loci, but rather are widespread with an enrichment at LADs on distal regions of chromosomes. Furthermore, small LADs appear to be gradually displaced from the NL by larger LADs. Additionally, we found that replicating DNA shows transiently increasing lamin contacts.

## Results

### Principle of pA-DamID to map and visualize NL associated DNA

The pA-DamID method is a hybrid of the CUT&RUN and DamID technologies (**Fig. 1A**). In CUT&RUN, cells are permeabilized and incubated with an antibody against a nuclear protein of interest, followed by protein A (pA) fused to micrococcal nuclease (MNase). Subsequent activation of the tethered MNase by Ca^++^ ions results in excision of DNA sequences that are in molecular proximity to the protein of interest. These sequences are then identified by high-throughput sequencing [25]. For pA-DamID, we used a highly similar strategy, except that we replaced the pA-MNase fusion protein by a pA-Dam fusion protein, which is then activated by addition of its methyldonor S-adenosyl-methionine (SAM). This results in adenine methylation (^m6^A) of DNA that is in molecular proximity of the protein of interest. The pattern of deposited ^m6^A can then be mapped genome-wide as in conventional DamID [24].

**Figure 1.**
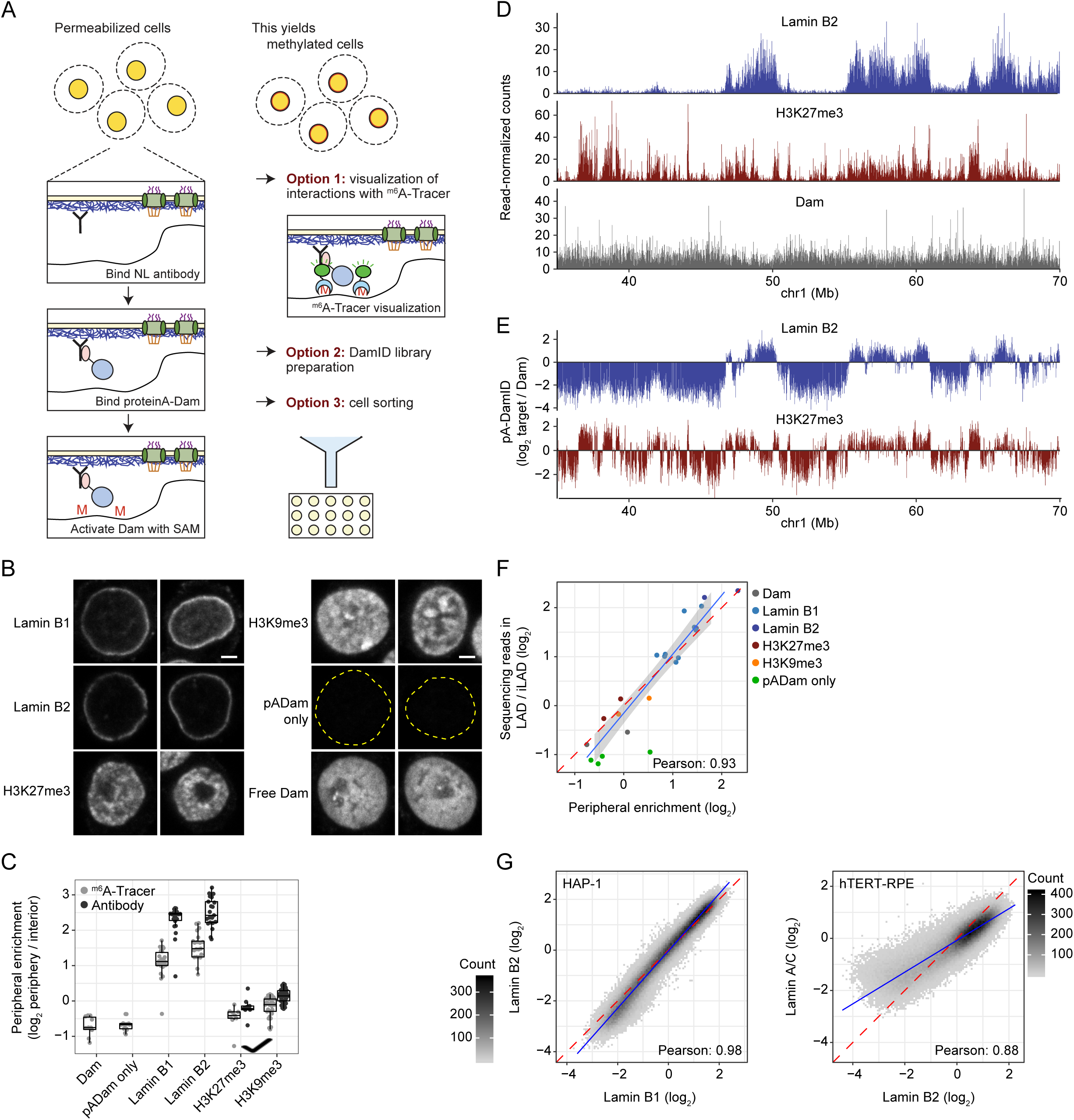
pA-DamID can visualize and map NL interactions. (**A**) Schematic overview of pA-DamID. Permeabilized cells are incubated with a primary antibody against a NL component, which in turn is bound by a pA-Dam fusion protein. Nearby DNA is then ^m6^A methylated upon Dam activation by incubation with SAM. The ^m6^A pattern can be visualized by staining with fluorescent ^m6^A-Tracer protein, or mapped genome-wide. Optionally, cells can be flow sorted to isolate specific cell populations prior to genome-wide mapping of ^m6^A. All steps are performed on ice, with the exception of 30-minute Dam activation at 37 °C. (**B**) Representative confocal microscopy sections of ^m6^A-Tracer signal in HAP-1 cells following pA-DamID with indicated antibodies. pA-Dam only: primary antibody was omitted, the dotted yellow line indicates DAPI segmentation. Free Dam: permeabilized cells were treated with freely diffusing pure Dam protein and SAM for 30 min. Scale bar corresponds to 2 µm. (**C**) Quantification of peripheral enrichment of antibody staining and ^m6^A-Tracer signals after pA-DamID with indicated antibodies. The nuclear rim and interior were segmented using DAPI signal and the mean ^m6^A or antibody signal was determined and tranformed to a log_2_-ratio. Every point represents a single cell; results are from one representative experiment. (**D**) Example of raw pA-DamID data tracks from 2 million HAP-1 cells for Lamin B2, H3K27me3 and the Dam control. Sequenced reads are counted in 20kb bins. (**E**) Same pA-DamID tracks of Lamin B2 and H3K27me3 as in (D) but after normalization to the Dam-only control (to correct for accessibility and amplification biases) and log_2_-transformation. (**F**) Correlation between the median peripheral enrichment of ^m6^A-Tracer determined by confocal microscopy and enrichment of sequencing reads within LADs in HAP-1 cells. LAD definition is based on conventional Lamin B1 DamID data. Every point represents a single pA-DamID experiment. (**G**) Comparisons of pA-DamID genome-wide data (bin size 20 kb) for different lamin antibodies: Lamin B1 vs. Lamin B2 in HAP-1 cells (*left panel*) and Lamin B2 vs. Lamin A/C in hTERT-RPE cells (*right panel*). Data are averages of at least two independent biological replicates.

An important advantage of pA-DamID is that the labeled DNA can also be visualized *in situ* using the ^m6^A-Tracer, before this DNA is sequenced. As will be shown below, this microscopy visualization can serve as a powerful quality control (e.g. to check that the tagged DNA is indeed close to the protein of interest) and can provide new biological insights. Additionally, labeled cells can be sorted following pA-DamID to study (rare) subpopulations of interest.

### Proof-of-principle of pA-DamID visualized by ^m6^A-Tracer staining

First, we expressed and purified pA-Dam fusion protein (**Fig. S1A**). We confirmed that the Dam moiety of the fusion protein was active, as judged from its ability to protect an unmethylated plasmid from digestion by the restriction enzyme Mbo I, which only cuts unmethylated GATC motifs (**Fig. S1B**). In addition, we expressed and purified ^m6^A-Tracer protein (**Fig. S1C**). Staining of fixed cells expressing either Dam or Dam-Lamin B1 with this protein showed the expected pan-nuclear and peripheral fluorescence pattern, respectively (**Fig. S1D**). We concluded that the purified pA-Dam and ^m6^A-Tracer proteins were functional.

We then applied the pA-DamID protocol (**Fig. 1A**) to human HAP-1 cells using antibodies against Lamin B1, Lamin B2 and the histone modifications H3K27me3 and H3K9me3. Visualization with purified ^m6^A-Tracer (**Fig. 1B**) and subsequent quantitative image analysis (**Fig. 1C**) showed that pA-DamID with both lamins yielded a clear rim staining. In contrast, the ^m6^A signals obtained with antibodies against H3K27me3 and H3K9me3 were found throughout the nucleus with only a modest enrichment at the periphery. A no-antibody control yielded virtually no staining, and permeabilized cells incubated with free Dam in the presence of SAM showed a homogenous nuclear staining except for the weakly labeled nucleoli (**Fig. 1B**). These data indicate that application of the pA-DamID protocol results in adenine methylation of DNA at the expected nuclear locations.

When Dam-Lamin B1 is expressed in vivo for 5-25 hours during interphase, LADs that interact with the NL become progressively labeled, eventually resulting in a layer of labeled chromatin of up to ∼1 µm thick [8]. This is because LADs are in dynamic contact with the NL. We expected that in pA-DamID this layer would be thinner, because the NL-tethered Dam is only activated for 30 minutes. In addition, permeabilization depletes small molecules including ATP and thus prevents active DNA remodeling in the nucleus [26]. Indeed, pA-DamID yields a ^m6^A layer that is ∼2.5 fold thinner than the layer in cells that express Dam-Lamin B1 *in vivo* (**Fig. S2A-C**). This is not an artifact due to collapse of chromatin onto the NL caused by the permeabilization, because permeabilization of cells expressing Dam-Lamin B1 *in vivo* did not significantly reduce the thickness of the ^m6^A layer compared to directly fixed cells (**Fig. S2C**). The thin layer of labeled DNA obtained by pA-DamID points to an improved temporal resolution of pA-DamID compared to conventional DamID.

### Genome-wide pA-DamID maps are reproducible and specific

Encouraged by these results, we proceeded to generate genome-wide pA-DamID maps, using a Lamin B2 antibody in HAP-1 cells (**Fig. 1D, top panel**). Amplification of ^m6^A-marked DNA fragments from genomic DNA was dependent on both the antibody and SAM (**Fig. S3A**), indicating that the mapping procedure is specific. For comparison, we also generated pA-DamID maps with an antibody against H3K27me3 (**Fig. 1D, middle panel**). Consistent with previous DamID studies [1, 27], Lamin B2 yielded a clear pattern of domains (LADs) that only partially overlaps with the H3K27me3 domains. Furthermore, incubation with freely diffusing Dam (instead of pA-Dam tethered by an antibody) resulted in a more homogeneous pattern that presumably reflects chromatin accessibility (**Fig. 1D, bottom panel**).

In conventional DamID, ^m6^A maps obtained with a Dam-fusion protein are typically normalized to a Dam-only control. This is done to correct for local variation in chromatin accessibility and for possible biases in PCR amplification or sequencing [28]. We applied a similar normalization in pA-DamID, using the data obtained with freely diffusing Dam as reference (**Fig. 1E**). Reproducibility between biological replicates was high (**Fig. S3B**). Noteworthy, the visualized peripheral enrichment of ^m6^A-Tracer signals correlates strongly with the enrichment of sequencing reads in LADs according to genome-wide DamID maps (**Fig. 1F**). Thus, the pA-DamID maps are generally consistent with the microscopy data.

Conventional DamID has indicated that Lamin B1, Lamin B2 and Lamin A/C all interact with the same LADs [29, 30]. Consistent with these results, we did not observe any notable differences between maps generated by pA-DamID with antibodies against these three lamins (**Fig. 1G**), although for Lamin A/C we observed a somewhat smaller dynamic range. This may be due to some interactions of Lamin A/C with inter-LAD regions [31, 32]. Together, these results show that pA-DamID can be used to map NL associations effectively.

### pA-DamID and DamID maps are highly similar but not identical

Next, we systematically compared DamID and pA-DamID in four different human cell types: HAP-1, HTC116, hTERT-RPE and K562. Overall, DamID and pA-DamID patterns are highly similar (**Fig. 2**, **Fig. S4**), although DamID data tended to have a larger dynamic range. In some cell types, especially in HCT116 and hTERT-RPE cells, we noted local discrepancies between the two methods (**Fig. 2A**, bottom panel). These differences involve mostly regions with low signals in DamID that have higher signals in pA-DamID. However, such differences are not obvious in HAP-1 and K562 cells.

**Figure 2.**
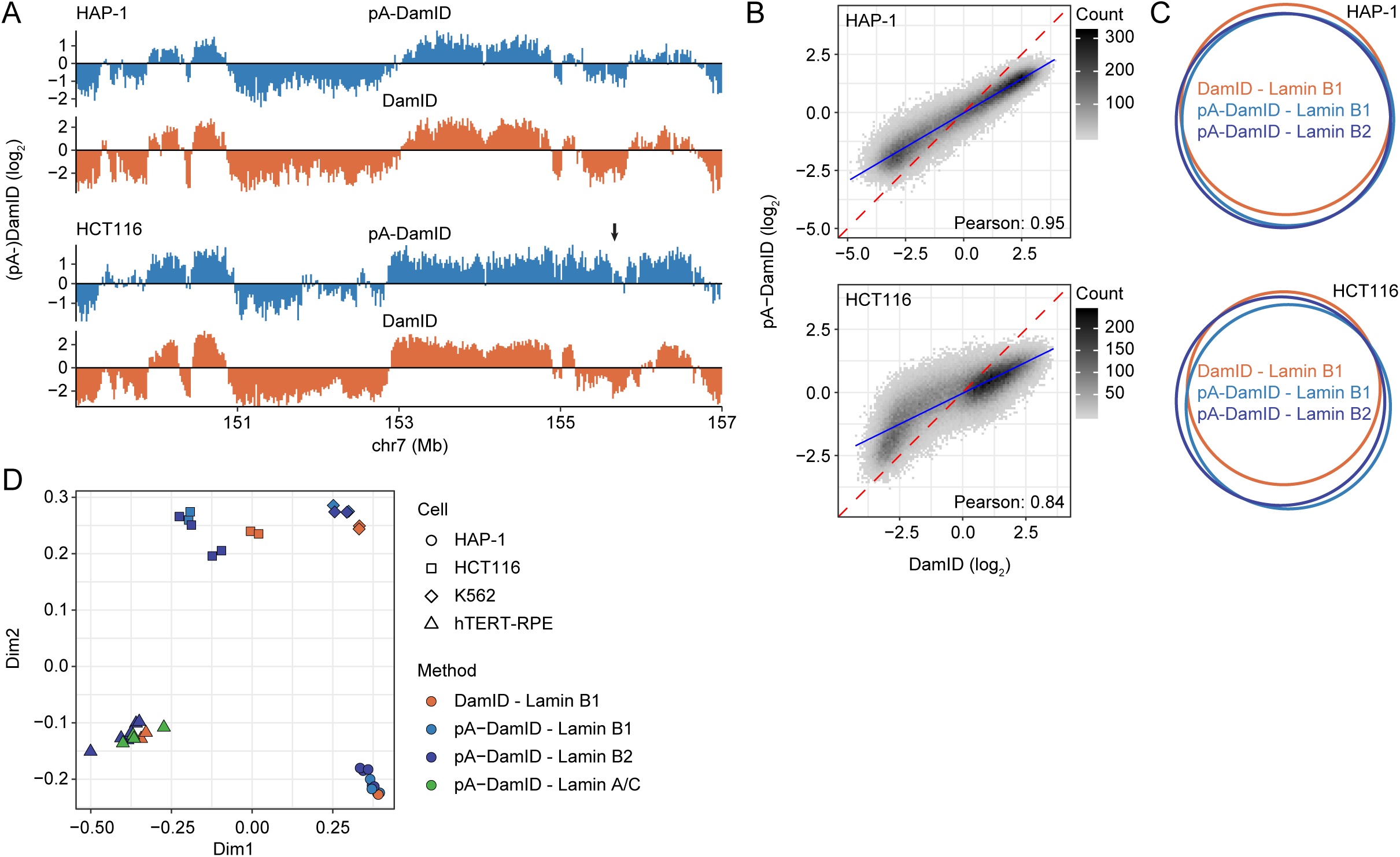
Comparison of NL interactions determined by pA-DamID and DamID. (**A**) Comparison of pA-DamID and DamID data across a representative genomic locus in HAP-1 and HCT116 cells. Arrow highlights a region in HCT116 cells with deviating signals between the two methods. (**B**) Genome-wide correlation of pA-DamID and DamID data. (**C**) Venn Diagram showing the overlap in LADs defined by a Hidden Markov Model. Results in (A-C) are based on averaged data from at least two independent experiments, bin size 20 kb. (**D**) Multidimensional scaling (MDS) plot [61] showing all individual replicates of DamID and pA-DamID experiments for various lamins and cell types.

We considered that the discrepancies could be explained by differences in nuclear roundness in the two assays. In DamID adherent cells remain attached to the culture dish, while in pA-DamID we normally harvest the cells by trypsinization before applying the protocol. Indeed, nuclei of hTERT-RPE cells are flat in adherent cells, and become less flat in trypsinized cells (**Fig. S5A-B**). Nevertheless, pA-DamID maps generated in cells that remained adherent (by leaving out the trypsinization step and performing the incubations in the culture dish) showed high correlation with those obtained after trypsinization (**Fig. S5C-D**). Thus, the change in nuclear shape does not affect NL interactions.

We emphasize that the differences between the DamID and pA-DamID maps are subtle compared to the differences between cell types, as is illustrated by dimensionality reduction analysis, which consistently clusters the datasets by cell type and not by method (**Fig. 2D**). As will be discussed below, the minor discrepancies between the two methods may arise from a difference in ^m6^A deposition during the cell cycle.

### NL interactions initiate throughout the genome in early G1

We then took advantage of the better time resolution of pA-DamID to study genome - NL interactions during the cell cycle. Initially, we explored how these interactions are established genome-wide after mitosis. To do so, we synchronized hTERT-RPE cells in prometaphase by sequential treatment with inhibitors of CDK1 (RO-3306) and microtubule polymerization (nocodazole), followed by mitotic shake-off. Counting of mitotic cells indicated that 90% and 93% of the cells were synchronized for the two replicates (**Fig. S6A**). We re-seeded these cells and mapped NL interactions by pA-DamID at various time points from 0 to 21 hours (**Fig. 3A**). These time points fall within one average cell cycle, which we estimated to take ∼26 hours (see Methods). At all time points except at 0 hours (i.e. when cells are in metaphase), ^m6^A-labeled DNA could be readily amplified (**Fig. S6B**). The lack of labeling in metaphase cells is consistent with lamins being depolymerized and excluded from the chromosomes at this stage [9, 10]. No genome-wide pA-DamID map could thus be obtained from this time point. The other time points yielded high-quality maps (**Fig. 3B**), although we observed modest differences in dynamic range of pA-DamID log-ratios (**Fig. S6C**, left panel). Because such differences were also present between biological replicates they are likely due to technical variation. To correct for this, we converted the log-ratios to z-scores (**Fig. S6C**, right panel).

**Figure 3.**
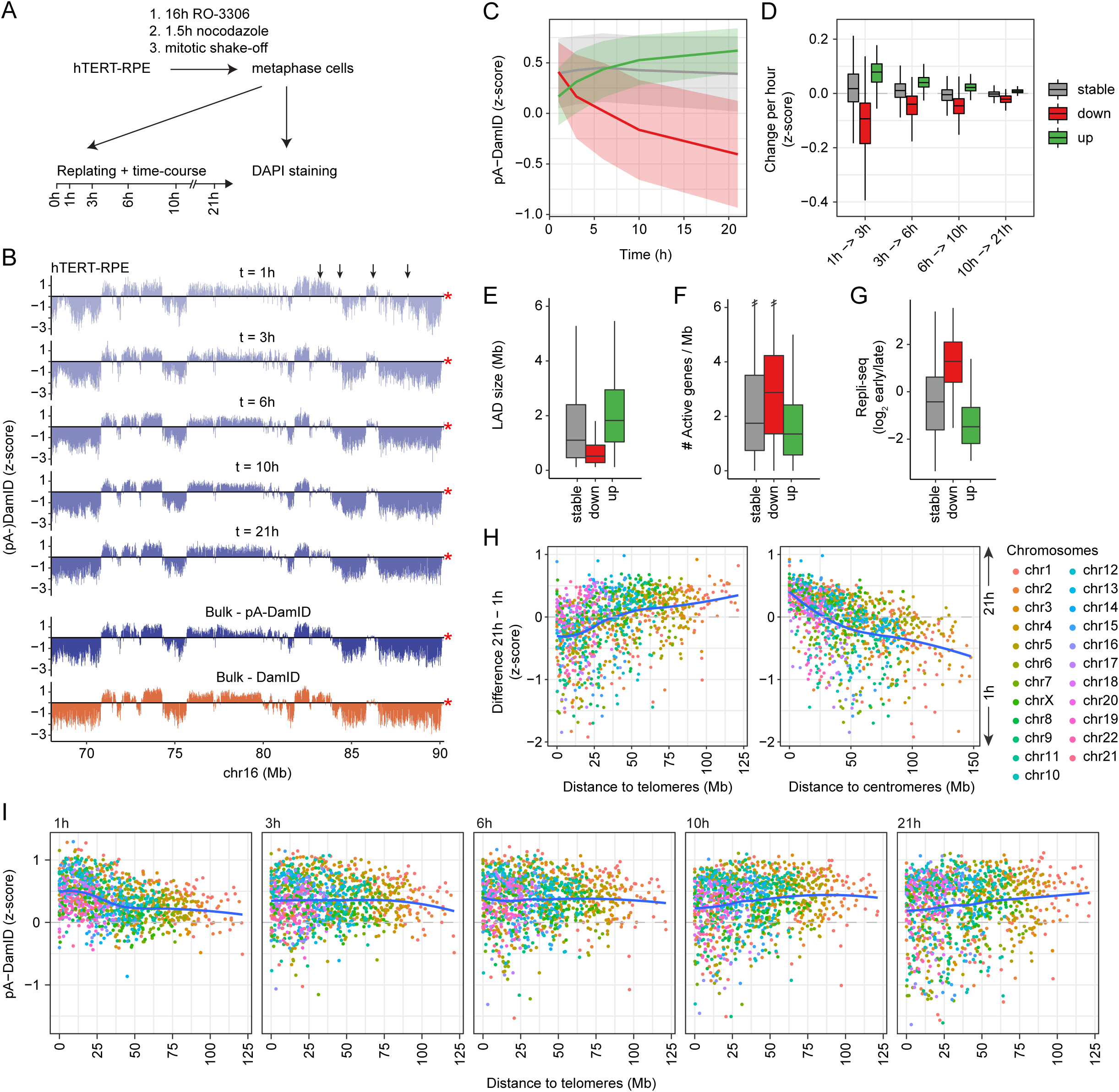
Mapping of NL interactions at various times after mitosis. (**A**) Strategy for synchronization of hTERT-RPE cells. Cells were first enriched in G2 phase by 16 hours incubation with 2.0 µg/mL RO-3306, washed, and incubated for another 1.5 hours with 25 ng/mL nocodazole to arrest cells in metaphase. Metaphase cells were then collected by mitotic shake-off. A small fraction was used to determine the proportion of mitotic cells by DAPI staining and confocal microscopy (**Fig. S6A**), while the remainder was replated for pA-DamID time series analysis. (**B**) NL interactions in hTERT-RPE cells at indicated times after mitosis, as determined by pA-DamID of Lamin B2. Log_2_-ratios were convered to z-scores to account for differences in dynamic range between replicates (**Fig. S6C**). Average of two independent biological replicates. Arrows point to regions with early NL interactions that decrease later in interphase. Red star marks chromosome end. (**C**) pA-DamID signal as a function of time after mitosis, for stable (grey) and dynamic LADs (red and green: decreasing and increasing NL interactions, respectively). Lines represent mean signals and the shaded area one standard deviation on both sides. (**D**) Rate of changes in NL interactions (pA-DamID z-score per hour) as function of time after mitosis. (**E-G)** LAD size distribution (**E**), density of active genes (**F**) and Repli-seq score distributions (**G**) for stable and dynamic LADs. (**H**) Difference in pA-DamID z-scores between 1h and 21h time points, plotted against the distance to telomeres (*top panel*) or centromeres (*bottom panel*). Each dot represents a single LAD. Chromosomes are colored and ordered by size. Blue line is the fitted loess curve. (**I**) Similar as (H), but showing pA-DamID z-scores for individual time points.

Interestingly, the mapping data show less defined bimodal distributions for early time points compared to later time points (**Fig. S6C**), suggesting that the separation of LADs and inter-LADs becomes progressively more pronounced after mitosis. Nevertheless, already at 1 hour we observe widespread domain patterns of NL interactions (**Fig. 3B**, **S6D**). Progression from prometaphase to late telophase in HeLa cells takes about 1 hour [33], suggesting that this timepoint captures the initial interactions with the reforming NL. Remarkably, the majority of these interactions is shared with later time points, indicating that most LADs can interact with the NL throughout interphase and are defined (and positioned at the NL) very soon after mitosis.

### Dynamic NL interactions correlate with chromosome location

Even though globally the pattern of interactions is similar throughout interphase, visual inspection suggested that some interactions become stronger over time, while others become weaker (**Fig. 3B**, **S6D**). To analyze this systematically, we defined a union set of 1176 LADs by a hidden Markov model per time point, and classified them either as static, increasing over time, or decreasing over time (**Fig. S6E**, Methods). Most LADs are stable, but 234 and 225 LADs show decreasing or increasing interactions over time, respectively (**Fig. 3C**). Changes occur mostly within the first few hours, although movements continue throughout interphase (**Fig. 3D**). Importantly, these dynamics are not related to variation in chromatin accessibility, because they are generally not accompanied by changes in Dam reads over time (**Fig. S6F**).

We next looked into characteristics of the dynamic LADs. At early time points, LADs with decreasing interactions do not have lower pA-DamID scores than stable LADs, suggesting that their detachment from the NL is not simply due to weak initial binding (**Fig. 3C**). However, decreasing LADs are generally smaller than stable and increasing LADs (**Fig. 3E**), which could make their interactions with the NL less persistent. In addition, decreasing LADs have a slightly higher density of active genes and replicate substantially earlier than stable or increasing LADs (**Fig. 3F-G**). Thus, decreasing LADs tend to carry euchromatin features that are untypical of LADs, which may explain why they detach from the NL within a few hours after mitosis.

Strikingly, we found that dynamic LADs have markedly skewed distributions along the chromosomes. Decreasing LADs are primarily located in distal regions of most chromosomes (typically within ∼25 Mb from the telomeres), and increasing LADs tend to be in centromere-proximal parts (**Fig. 3H, Fig. S6D**). Early after mitosis, distal LADs tend to have stronger interactions with the NL than proximal LADs (**Fig. 3I, S6G**). Over time this skew is inverted, with centromere-proximal LADs gradually gaining interactions and distal LADs showing weaker interactions. This dependency on chromosomal location is not linked to LAD size, because these features are not correlated (**Fig. S6H**; Pearson correlation: 0.01). These data indicate that chromosomes are gradually reoriented during interphase, with proximal LADs progressively taking over some of the NL interactions from distal LADs.

### LAD dynamics are linked to telomere distance and LAD size in multiple cell types

We considered that the observed dynamics may be artefacts of the drug treatments that we applied to obtain synchronized cells. In addition, the observations might be specific for hTERT-RPE cells. To address these points, we performed pA-DamID on unsynchronized populations of HAP-1, K562 and HCT116 cells, and then sorted cells based on DNA content to obtain G1, mid-S and G2 pools (**Fig. S7A-B**). Overall, the resulting maps are very similar between cell cycle stages but not identical (**Fig. 4A**, **S7C-D**).

**Figure 4.**
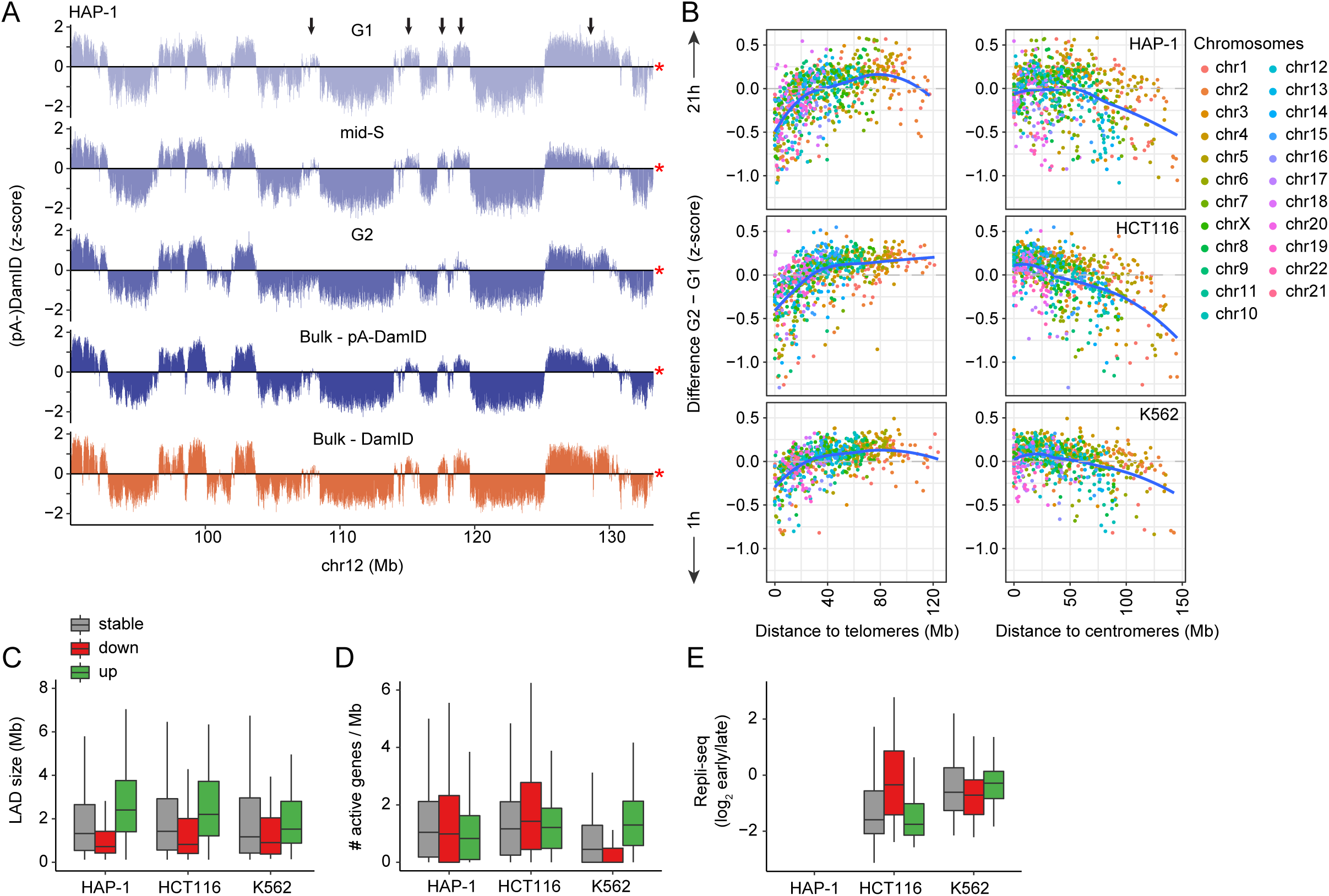
LAD dynamics in other cell types. (**A**) pA-DamID z-scores of a representative locus in HAP-1 cells sorted for G1, mid-S and G2 phases. Arrows point to regions that lose NL interaction from G1 to G2. (**B**) Similar plot as Fig. 3H, showing the difference in pA-DamID z-scores between G2 and G1 sorted pools as function of distance to either telomere or centromere for HAP-1, HCT116 and K562 cells. (**C-E**) LAD size distribution (**C**), density of active genes (**D**) and Repli-seq scores (**E**) for stable and dynamic LADs. Repli-seq data are not available for HAP-1 cells.

We separated LADs into three classes of stable, increasing or decreasing NL interactions over time by comparing the pA-DamID signals in G1 and G2 phases (**Fig. S7E**). Similar to what we observed in synchronized hTERT-RPE cells, decreasing interactions involved predominantly distal LADs (**Fig. 4B**, left panel). We also find that small LAD size is a consistent characteristic of decreasing interactions, although this is less pronounced in K562 cells (**Fig. 4C**). In contrast, increasing interactions for centromere-proximal LADs and enriched euchromatin features in decreasing LADs are only apparent for HCT116 cells (**Fig. 4B**, right panel, **Fig. 4D-E**). We conclude that the links between dynamic NL interactions and the chromosomal positions and sizes of LADs are not specific to synchronized hTERT-RPE cells, but apply more generally to cultured cells of various origins.

### S-phase chromatin shows increased lamin interactions

Next, we inspected the pA-DamID profiles obtained in mid-S phase in more detail. We frequently observed that LAD border regions or weak LADs have transiently increased pA-DamID scores in mid-S compared to G1 and G2 (**Fig. 5A-B**, **S8A-B**). These regions often coincide with areas that replicate in mid-S phase (**Fig. 5A**). To test this genome-wide, we calculated the changes in pA-DamID scores between mid-S and either G1 or G2, and plotted this against replication timing [34, 35] (**Fig. 5C**). Indeed, regions that replicate in mid-S phase show on average a small but consistent increase in NL interactions in mid-S cells compared to G1 and G2 cells. The increase in pA-DamID signal is not caused by a change in accessibility, as Dam reads remain constant except for DNA duplication of early replicating DNA (**Fig. S8C**).

**Figure 5.**
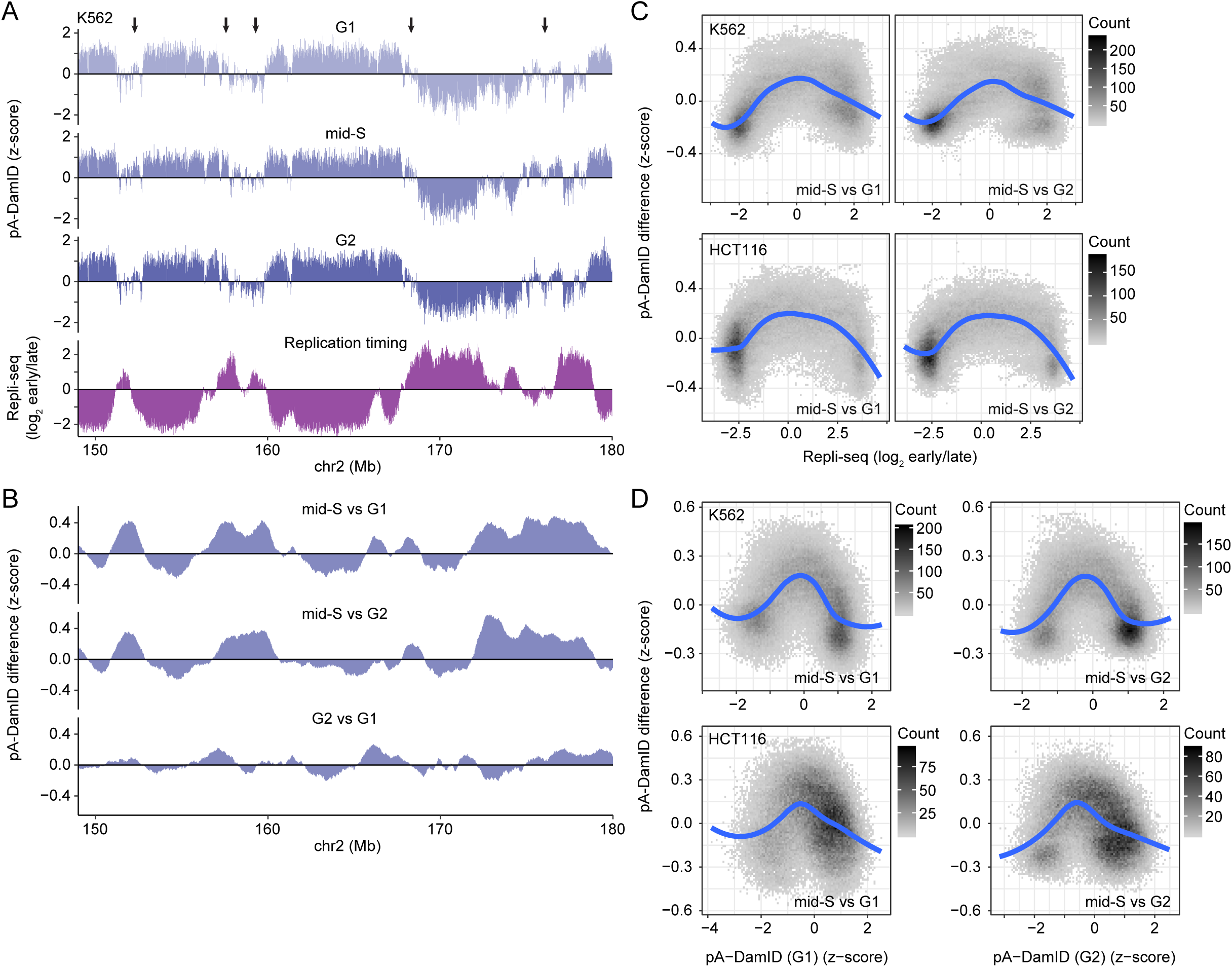
Mid-S chromatin has transiently increased Lamin B2 interactions. (**A**) Representative K562 locus comparing cell cycle sorted cells with replication timing. Arrows point to regions with increased mid-S signal compared to G1 and G2, which tend to overlap with regions that exhibit intermediate replication timing. (**B**) Difference tracks in pA-DamID z-scores between the cell cycle stages for the same locus as (A). Data were smoothed by a running mean of 5 bins (bin size 20 kb). (**C**) Scatterplots of Repli-seq vs. the difference in smoothed pA-DamID z-scores between mid-S sorted cells and either G1 or G2 sorted cells. Bins with pA-DamID differences between G1 and G2 bigger than 0.2 were filtered out. Blue line shows a fitted loess curve. (**D**) Scatterplots of changes in pA-DamID signal versus the pA-DamID signal in G1 (left-hand panels) or G2 cells (right-hand panels). Blue line shows a fitted loess curve.

We investigated whether the LADs with increased lamin interactions in S-phase were already partially associated with the NL in G1 phase. Indeed, the strongest increases in NL contacts during S-phase involve regions with intermediate pA-DamID signals in G1 (**Fig. 5D**, left-hand panels). In G2 phase, these regions revert again to intermediate pA-DamID signals (**Fig. 5D**, right-hand panels). These results indicate that S-phase chromatin, at least during mid-S phase, exhibits temporarily increased contacts with lamin B2, suggesting a transient strengthening of NL interactions.

### Minor differences between pA-DamID and DamID are linked to cell cycle bias

Finally, we revisited the minor differences between pA-DamID and DamID data in unsynchronized cells (**Fig. 2**). We asked whether DamID data might be somewhat skewed due to an unequal representation of specific stages of the cell cycle. For example, DNA replication converts DNA that was fully Dam-methylated in G1 into hemi-methylated DNA. The latter shows impaired cutting by DpnI, the restriction enzyme that is used to identify ^m6^A-methylated DNA in the DamID protocol [36]. Thus, NL interactions that occur in G1 phase may be somewhat under-detected by DamID in unsynchronized cells. To test this, we compared conventional DamID data to the pA-DamID data from each cell cycle phase. This indicated that the DamID data are most similar to G2 phase pA-DamID, especially for the HCT116 cells that showed the biggest differences (**Fig. S9**). This result suggests that the minor differences between pA-DamID and DamID in unsynchronized cells are mostly related to somewhat different representations of cells across the cell cycle.

## Discussion

In the past decade, DamID has been used to identify and characterize LADs [1, 4, 5, 22]. However, conventional DamID lacks the temporal resolution needed to study dynamics of LAD - NL interactions that occur within time spans of about one hour. Here, we present pA-DamID, a method to take “snapshots” of NL contacts. We applied pA-DamID to reveal the dynamics of LAD – NL contacts during initial interphase and during DNA replication.

### Cell cycle dynamics of genome – NL interactions

Our data indicate that most changes in NL contacts occur in the first hours after mitosis, consistent with the higher mobility of chromatin at this stage, as observed by microscopy [16]. Initial NL contacts are widespread throughout the genome and quickly mature in a more bimodal distribution. This is in agreement with single-cell Hi-C analysis in mouse embryonic stem cells, which showed that the B compartment (largely overlapping with LADs [4]) gradually moves towards a more radial positioning in early G1 [37]. Similarly, Hi-C in synchronized cells showed that chromatin compartments appear in telophase and gradually gain in strength [38, 39].

Earlier work indicated that telomeres are enriched near the nuclear periphery during postmitotic nuclear assembly [13]. Our results show that this enrichment is not limited to the telomeres, but rather involves distal chromosome regions up to ∼25 Mb from the telomeres, typically including multiple LADs. Interestingly, LAP2α, which can interact with Lamin A/C, has been found to localize on the distal regions of mitotic chromosomes, i.e. prior to NL assembly. Later, LAP2α migrates towards the nuclear interior [12]. Possibly, LAP2α marks mitotic chromosomal regions that are destined to become the early NL assembly sites in telophase. However, it is also possible that distal chromosome regions are more probable sites for early NL interactions because they are more likely to be at the surface of the telophase chromatin mass, as a consequence of the pulling motion on centromeres earlier in mitosis [12]. Because most LADs begin to interact with the NL soon after mitosis, it is possible that LADs are generally “bookmarked” for such interactions, for example by heterochromatin marks such as H3K9me2 [40].

*In situ* labeling experiments have shown that DNA replication occurs in a specific spatial sequence inside the nucleus, initiating in interior foci before moving towards the NL and nucleolus and ending in large nuclear foci corresponding to pericentric heterochromatin [41, 42]. According to these studies, juxtaposition of replicating DNA to the NL occurs primarily during mid-S phase. In addition, B-type lamins have been found to overlap with replication foci in the nuclear interior during mid-late S phase [20]. Our pA-DamID results showing a transient increase in lamin B2 contacts with mid-S chromatin, are in agreement with both findings. The precise function of this transient association of lamins with S-phase chromatin remains to be elucidated.

### pA-DamID as a tool for studying NL interactions

pA-DamID is an alternative implementation of protein A-based enzyme targeting to study protein-DNA interactions, following the principle of CUT&RUN [25] and its derivative CUT&TAG [43]. Compared to CUT&RUN, a major advantage of pA-DamID is that ^m6^A-marked DNA can be visualized microscopically (using purified ^m6^A-Tracer). This provides a simple visual check that ^m6^A is deposited near the antibody used, in this case near the NL. Furthermore, in our pA-DamID protocol the genome-wide maps obtained with Dam tethered to a lamin antibody are normalized to maps obtained with untethered Dam. This way the data are corrected for local differences in chromatin accessibility, similar to the standard DamID protocol [24, 28]. We believe that such a normalization is important to reduce mapping biases. This may be particularly relevant for LADs, which are mostly heterochromatic. This normalization is to our knowledge not commonly applied in current CUT&RUN or CUT&TAG protocols [25, 43].

pA-DamID is inherently limited in resolution due to its dependence on GATC sequence motifs, but in contrast to CUT&RUN the template is not destroyed [25]. Processed cells can therefore be stained and sorted for specific subpopulations, as we illustrated here by flow sorting based on DNA content. Possibly this is also feasible with CUT&TAG, although the applicability for the NL has not been tested to our knowledge [43].

Similar to CUT&RUN and CUT&TAG, pA-DamID requires permeabilization of unfixed cells. This could potentially affect protein localization and genome organization. Upon digitonin treatment, most nuclear protein is retained but small amounts can leak out [44]. High concentrations of digitonin (>500 μg/mL) result in nuclear shrinking [45], but more subtle effects on genome architecture have not been ruled out at lower concentrations. However, by direct visualization of ^m6^A-tagged DNA we observed no effect of this permeabilization on the peripheral genome organization. Presumably, small molecules including ATP are depleted from the nucleus after permeabilization (as illustrated by the depletion of endogenous SAM in pA-DamID and Ca^++^ in CUT&RUN [25] and previous observations [26]). This may prevent active remodeling of the nuclear organization, enabling “snapshot” genome-wide mapping.

Compared to conventional DamID, pA-DamID yields data of approximately similar quality as DamID, although it has a slightly reduced dynamic range. This may be due to imperfect antibody-based localization of Dam. We also noticed a nucleosomal-like pattern of ^m6^A-labeled DNA fragments after pA-DamID (**Figure S3A**), which suggests an inability of Dam to methylate nucleosomal DNA in permeablized nuclei. This may reduce the number of GATC motifs available for ^m6^A tagging. A nucleosomal pattern has not been observed in conventional DamID [24], in which case ^m6^A is deposited in living cells, where nucleosomes may undergo a more dynamic repositioning than in permeabilized cells. A major practical advantage of pA-DamID is that it does not require *in vivo* expression of a Dam-fusion protein. This will greatly facilitate the mapping of NL contacts in primary cells and tissues. Together with the increased time resolution, this creates new opportunities to study the role of the NL in genome organization [46].

## Acknowledgements

We thank the NKI Genomics, Flow Cytometry, Protein, Digital Microscopy and RHPC core facilities for technical assistance. We thank Andrew Belmont, Jian Ma, David Gilbert and other members of the 4DN Center for Nuclear Cytomics for helpful discussions. We thank the lab of David Gilbert for sharing Repli-seq data prior to publication. We thank the lab of Jop Kind for sharing a new HT1080 inducible Dam-Lamin B1 clone. Supported by NIH Common Fund “4D Nucleome” Program grant U54DK107965 (BvS). The Oncode Institute is partly supported by KWF Dutch Cancer Society.

## Author contributions

TvS: Conceived and designed study, conducted majority of experiments and data analysis, wrote manuscript. DPH: Performed experiments. MV: Performed experiments. BvS: Designed study, wrote manuscript, supervised project.

## Methods

### Cloning

The DNA sequence encoding the proteinA-Dam fusion protein was codon optimized for bacterial expression and ordered as gBlock (IDT). Sequences for ligation-idependent cloning (LIC) were added to the flanks and used to insert the gBlock into the pETNKI-his-3C-LIC-kan vector [47]. ^m6^A-Tracer DNA (GFP-DpnI*) was PCR amplified from the original plasmid [8] with LIC sequences attached to the primers. The PCR fragment was purified and cloned by LIC into the pFastBacNKI-his-3C-LIC vector (Luna-Vargas, 2011, 21453775}. We also replaced GFP with a HALO tag in the ^m6^A-Tracer plasmid with PCR amplification and cloning into the AgeI and BglII restriction sites. This plasmid was processed identical to the GFP plasmid to create an expression vector and to obtain purified protein.

### Expression and purification of his-pA-Dam

The his-pA-Dam fusion protein was expressed in Rosetta2(DE3) cells in LB medium supplemented with 30 µg/ml kanamycin and 40 µg/ml chloramphenicol. Cells were grown at 37°C to OD_600_ = 0.6-0.8. Cells were cooled to 20°C before 0.4 mM IPTG was added and protein was expressed overnight. Cells were harvested by centrifugation (1 minute 3,000g) and resuspended in lysis buffer (25 mM Tris pH 8.0, 200 mM NaCl, 1 mM TCEP). Cells were lysed by sonication and the lysate was clarified by centrifugation (30 minutes 50,000g at 4 °C). The soluble fraction was loaded on 1 mL Nickel beads (GE Healthcare). Beads were washed with lysis buffer containing 20 mM imidazole before protein was eluted by 250 mM imidazol in lysis buffer. Fractions were analysed by SDS-PAGE, pooled and concentrated. Protein was diluted 4-fold with 25 mM Tris pH 8.0 and purified by anion-exchange chromatography using a Resource Q column (GE Healthcare), equilibrated in 25 mM Tris pH 8.0, 1 mM TCEP. Protein was eluted by a NaCl gradient (20 – 500 mM) in the same buffer. Fractions were concentrated and glycerol was added to a final concentration of 50 %. His-pA-Dam protein was stored at -20 °C. About 0.1 mg of pA-Dam could be purified from 1 liter of Rosetta2(DE3) culture.

### Expression and purification of his-^m6^A-Tracer

The pFastBacNKI-his-3C-^m6^A-Tracer construct was used for creation of baculovirus according to the Bac-to-Bac system (Thermo Fisher Scientific). Briefly, plasmid DNA was transformed into *E. coli* EMBACY cells [48] and positive clones were selected by blue/white screening. Bacmid DNA was isolated and used for transfection of sf9 insect cells. 10 µg of DNA was incubated with 5 µl of CellFectin reagent (Thermo Fisher Scientific) and added to sf9 cells grown in Insect-Express medium (Lonza) in 6-well plates (Corning). After 3 days, medium containing the virus (P0 virus) was harvested and added to 50 ml of sf9 suspension culture. After 3 days of infection, medium containing P1 virus was harvested and after addition of 0.5% FCS, P1 virus was stored in the dark at 4 °C. 500 mL of sf9 cells (density 1 million cells/mL) were infected with 500 µl P1 virus and his-^m6^A-Tracer was expressed for 3 days. Cells were harvested by centrifugation (15 minutes, 1,000 g) and resuspended in lysis buffer (25 mM Tris pH 8.0, 100 mM NaCl, 0.5 mM TCEP). After sonication and centrifugation (30 minutes, 50,000 g), the soluble lysate was loaded onto 1 mL Nickel beads (GE healthcare). The column was washed with lysis buffer containing 15 mM imidazole and his-^m6^A-Tracer was eluted by 250 mM imidazole in lysis buffer. Fractions were pooled and diluted 1:1 with milliQ water and loaded onto a 6 ml Resource S column. His-^m6^A-Tracer was eluted by a NaCl gradient (20 to 500 mM) in 25 mM Tris, 0.5 mM TCEP. Fractions were pooled and loaded on a SEC70 size exclusion chromatography column (Bio-Rad), equilibrated with 25 mM Tris pH 8.0, 150 mM NaCl, 0.5 mm TCEP. Fractions were pooled, concentrated and flash frozen in liquid nitrogen before storing at -80 °C. A working batch was supplemented with 50% glycerol and stored at -20 °C. About 1.5 mg of his-^m6^A-Tracer could be purified from 1 liter of sf9 insect cell culture. A HALO-tagged ^m6^A-Tracer was produced using the same strategy.

### MboI protection assay

Dam activity of the purified pA-Dam proteins was assayed as follows. Unmethylated control plasmid was produced in *dam^−^* bacteria (New England BioLabs #C2925H) and methylation status was confirmed by DpnI and DpnII digestion. A concentration range (0.125, 0.5, 2 µL) of pA-Dam protein was incubated with 500 ng of unmethylated plasmid in 20 µL of 1x dam MethylTransferase buffer (New England BioLabs #M0222S) supplemented with 80 µM S-adenosylmethionine (SAM) for 30 minutes at 37°C. As a positive control, a concentration range (0.25, 1, 4, 16 units) of Dam enzyme was used (New England BioLabs #M0222S). Next, the reaction mix was heat inactivated for 15 minutes at 65°C, and 10 µl was mixed with 40 µL of 1x NEBuffer 3 supplemented with 10 mM MgCl_2_ and 5 units of Mbo I (New England BioLabs #R0147L). This reaction was incubated for 1 hour at 37°C, followed by analysis on agarose gel. For Dam activity, one unit is defined by New England Biolabs as the amount of enzyme required to protect 1 µg (*dam*-) Lambda DNA in 1 hour at 37°C in a total reaction volume of 10 µl against cleavage by MboI restriction endonuclease.

### Cell Culture

HAP-1 cells were cultured in IMDM (Gibco) supplemented with 10% fetal bovine serum (FBS). K562 cells (ATCC CCL-243), HCT116 cells (ATCC CCL-247) and hTERT-RPE cells (ATCC CRL-4000) were cultured according to ATCC protocol. For HCT116, 0.05% Trypsin-EDTA (Gibco) was used for passaging according to 4D Nucleome guidelines (https://data.4dnucleome.org/biosources/4DNSRMYUIVGD/). Cells were tested for mycoplasma every 2-3 months.

Cell cycle duration of hTERT-RPE cells was estimated by passaging the cells and counting the number of seeded and harvested cells over a period of 24 and 48 hours (Bio-Rad, TC20).

HT1080 cells expressing inducible Dam-Lamin B1 (new clone kindly provided by J. Kind [8]) and wildtype HT1080 cells were cultured in DMEM supplemented with 10% FBS and 1% penicillin and streptomycin. Dam-Lamin B1 was stabilized for 16 hours with 500 nM Shield1.

### Cell synchronization

hTERT-RPE cells were synchronized in mitosis (metaphase) using a sequence of G2 enrichment, mitotic synchronization and shake-off. First, cells were cultured for 16 hours with 2.0 μg/mL RO-3306 (CDK1 inhibitor) to enrich cells in G2. Cells were then washed three times with DMEM-F12 and incubated for another 1.5 hours with 25 ng/mL nocodazole (microtubule inhibitor) to synchronize in metaphase. This resulted in ∼40% mitotic cells estimated by morphology. Next, mitotic shake-off was used to obtain a nearly pure population of synchronized metaphase cells. The majority of synchronized cells was re-plated and used to determine NL contacts during interphase, but a fraction was used to estimate synchronization efficiency. For the latter, DigWash-permeabilized cells (see pA-DamID below) were put on poly-L-lysine coated coverslips (see below) and fixed for 10 minutes in 2% formaldehyde in phosphate-buffered saline (PBS). After washes with PBS and H_2_O, coverslips were mounted with Vectashield + DAPI (Vector Laboratories, #H-1200), dried and sealed with nail polish. The percentage of mitotic cells was determined manually by counting metaphase and anaphase appearance from confocal sections. For the replated cells, light microscopy analysis confirmed adherence of the cells to the culture dish and completion of cell division.

### Lamin B1 DamID

DamID-seq and data processing were performed as reported previously [49]. The K562 DamID data are also from [49]. To prevent amplification of apoptotic fragments present in HAP-1 cell culture, DNA was first dephosphorylated before DpnI digestion. To do so, up to 500 ng of gDNA was incubated in 10 µL H_2_O with 1x CutSmart buffer and 0.5 U rSAP (New England BioLabs #M0371L) for 1 hour at 37°C, followed by heat inactivation for 10 minutes at 65°C. Next, 10 U DpnI (New England BioLabs #R0176L) was added and sample processing continued as described [49].

DamID and pA-DamID sequence reads are binned in 20 kb bins, which provides a good balance between resolution and data quality for the synchronization and cell sorting experiments. For data visualization, the mean signal of all replicates was used.

### pA-DamID

A detailed protocol of pA-DamID is provided as **Supplementary Information S1.** pA-DamID is a hybrid of DamID and CUT&RUN [25]. It is performed with similar buffers as CUT&RUN for consistency, although there is no need to deplete bivalent ions in pA-DamID. One million cells were harvested by centrifugation (3 minutes, 500 g, this speed and duration were used in all subsequent wash steps) and washed in ice-cold phosphase-buffered salaine (PBS). Adherent cells were trypsinized before harvesting. For haploid HAP-1 cells, two million cells were used. Cells were washed with ice-cold digitonin wash buffer (DigWash) (20 mM HEPES-KOH pH 7.5, 150 mM NaCl, 0.5 mM spermidine (Sigma, #S0266-5G), 0.02% digitonin (Millipore, #300410-250MG), cOmplete Protease Inhibitor Cocktail EDTA-free (Roche, #11873580001)). Permeabilized cells were resuspended in 200 µL DigWash with a primary antibody and slowly rotated for 2 hours at 4°C, followed by a wash step with 0.5 mL DigWash buffer. In case of mouse or goat primary antibodies, the above steps were repeated with a secondary rabbit antibody for 1 hour. Nuclei were resuspended in 200 µL DigWash with 1:100 pA-Dam (between 20-60 New England BioLabs units, **Fig. S1B**) and rotated for 1 hour at 4°C, followed by 2 washes with 0.5 mL DigWash to remove unbound pA-Dam. Dam was activated by resuspension in 100 µL DigWash supplemented with 80 µM SAM (New England BioLabs, #B9003S) and incubated for 30 minutes at 37°C. The reaction was stopped by cooling the samples to 4°C and washing with 0.5 mL DigWash. Library preparation for high-throughput sequencing is identical to conventional DamID [24, 49], except that we left out the DpnII digestion as this digestion step prevented almost all amplification. Possibly, this is because methylation by pA-Dam *in vitro* is sparser than by expression of Dam proteins *in vivo*, and the DpnII digestion suppresses amplification unless two neighboring GATC motifs are both methylated.

We routinely included three controls: First, a negative control without any Dam enzyme. This control should be completely negative for ^m6^A. Second, a pA-DamID sample without a primary antibody, which would reveal potential non-specific binding of pA-Dam. Third and most important, a sample without antibody or pA-Dam incubation, but with 0.5 µL Dam enzyme (New England BioLabs #M0222S) added during the activation step. This control is used to normalize pA-DamID signals for local accessibility and amplification bias.

pA-DamID on-the-plate was performed identical to regular pA-DamID, but with cells still attached in a 6-well plate. The volumes were increased to 400 µL for every step to fully cover the cells, and plates were slowly shaken instead of rotated.

For pA-DamID we generally used primary antibody dilutions similar to those used for immunofluorescence labling. We used a 1:100 dilution for Lamin B2 (Abcam ab8983, mouse), Lamin A/C (SCBT sc-6215, goat), H3K27me3 (CST C36B11, rabbit) and H3K9me3 (Abcam ab8898, rabbit). A 1:500 dilution was used for Lamin B1 (Abcam ab16048, rabbit). For mouse primary antibodies, a secondary rabbit anti-mouse was used in 1:100 dilution (Abcam ab6709, rabbit). For goat antibodies we used rabbit anti-goat (ab6697) diluted 1:100.

### Cell cycle sorted pA-DamID

For pA-DamID of cells in specific cell-cycle phases, pA-DamID was first performed as described above for HAP-1, K562 and HCT116 cells, using the Lamin B2 antibody as well as a Dam-only control as described above. Next, the cells were washed once in PBS with 2% fetal bovine serum, and resuspended in the same buffer with 2 µg/mL propidium iodide (PI). Cells were sorted by PI signal in a MoFlo Astrios cell sorter (Beckman Coulter), with gate settings as shown in **Fig. S7A-B**. Sorted cells were collected in a 96-well PCR plate with each well pre-filled with 3 µL 1.33x lysis buffer (1x lysis buffer: 10 mM Tris acetate pH 7.5, 10 mM Mg acetate, 50 mM potassium acetate, 0.67% Tween 20, 0.67% Igepal, 0.67 mg/ml Proteinase K). For every sample and cell cycle phase, 2 or 3 wells were filled with 1000 cells, corresponding roughly to 1 µL. Proteinase K digestion was performed for 4 hours of 53°C and was heat inactivated for 10 minutes at 80°C. Further processing was done according to a previously reported single-cell DamID protocol [50], using 22 cycles of PCR amplification.

### ^m6^A-Tracer visualization of pA-DamID cells

Nuclear staining with ^m6^A-Tracer was performed before fixation to minimize background signal. Coverslips were first coated with 0.1% (w/v) poly-L-lysine (Sigma-Aldrich, #P8920) for 15 minutes, washed with H_2_O (1x) and PBS (3x) and stored in 70% ethanol for later use. Processed cells were centrifuged (3 minutes, 500 g) and resuspended in 100 µL DigWash (see pA-DamID section) or PBS with 1:500 ^m6^A-Tracer protein (1.15 mg/mL) and 1:500 secondary anti-rabbit antibody (Jackson 711-585-152, donkey, Alexa 594) and rotated for 1 hour at 4 °C. Cells were washed 2x with 0.5 mL cold DigWash or PBS to remove unbound fluorophores and then bound to poly-L-lysine coated coverslips on ice. The last wash was always performed with PBS. Cells were fixed at room temperature on the coverslips for 10 minutes with 2% formaldehyde/PBS and washed with PBS. After a final wash with H_2_O, coverslips were mounted with Vectashield + DAPI (Vector Laboratories, #H-1200), dried and sealed with nail polish.

### Effect of pA-DamID on endogenous LADs

HT1080 cells expressing inducible Dam-Lamin were plated both on coverslips and in a regular culture dish and Dam-Lamin B1 expression was activated overnight with 500 nM Shield1 (Aobious #AOB1848). Cells grown in the culture dish were trypsinized and processed with the pA-DamID protocol, but without pA-Dam or Dam added. The cells were then fixed on poly-L-lysine coated coverslips as described above. As soon as the culture dish cells were permeabilized, the cells on coverslips were fixed with 2% formaldehyde/PBS for 10 minutes, permeabilized for 20 minutes with 0.5% NP-40/PBS and blocked with 1% BSA/PBS for 1 hour. Next, coverslips were incubated for 1 hour at room temperature with 1:500 Lamin B1 antibody (Abcam ab16048, rabbit) and washed 3x with PBS. PBS with 1:500 ^m6^A-Tracer protein (1.15 mg/mL) and 1:500 secondary anti-rabbit antibody (Jackson 711-585-152, donkey, Alexa 594) was added and incubated for 1 hour, followed by washing with PBS (3x) and H_2_O (1x), and mounted with Vectashield + DAPI. For comparison, wildtype HT1080 cells (not expressing Dam-Lamin B1) were subjected to pA-DamID using Lamin B2 antibody and processed as the HT1080 Dam-Lamin B1 HTC75 cells.

### Microscopy and image analysis

Single 1024×1024 confocal sections were imaged on a Leica SP5 with a 63x NA 1.47 oil immersion objective, using bidirectional scanning, 3x electronic zoom and 8x line averaging. Laser power varied between experiments but was kept constant within each experiment.

Image analysis was performed in ImageJ 2.0.0 and R. To calculate peripheral enrichment, nuclei were segmented on the DAPI staining and split into a peripheral ring and interior. Mean signal was determine for both compartments. Peripheral enrichment was defined as the log_2_ ratio of the mean peripheral compartment signal over the interior compartment. The peripheral compartment was defined as the outline of a DAPI mask dilated by 2 pixels to capture all signal. This mask thus extended the boundaries of the nucleus and could lead to a negative enrichment.

To calculate halfway decay distances, DAPI-segmented nuclei were morphologically dilated by 2 pixels and a distance map to the mask edge was calculated. This distance map was used to determine the mean pixel-score to the periphery for lamin B2 antibody and ^m6^A-Tracer. A trimmed-mean was used to make the average more robust to outliers, where 20% on both sides was discarded. For both the lamin B2 antibody and ^m6^A-Tracer, an exponential decay function was fitted starting from the maximum value and used to determine the halfway decay distance. The difference in halfway decay distances was used as measure of LAD positioning relative to the NL.

### Differential analysis for LADs

We accounted for differences in dynamic range in pA-DamID signals between experiments by converting the log_2_ (Lamin B2:Dam) data to z-scores (mean of zero and a standard deviation of one), thus keeping the data distribution identical. For every population (i.e. HCT116 G1; hTERT-RPE 3hr), a consensus LAD definition was determined by selecting LADs that are called in more than half of the replicates using hidden Markov modeling (https://github.com/gui11aume/HMMt). Next, we determined the union across different conditions (i.e. HCT116 G1 and G2). The LAD score was defined as the mean signal of scaled data tracks.

Significant changes were called using a modified limma-voom approach [51]. Limma-voom is a limma extension that models the mean-variance relationship of log-counts. The reasoning is that high counts generally have lower variation. DamID data is already in log scale and can easily be adapted for this purpose. However, a high log-ratio in DamID can be supported by few reads and thus have a high variation. Instead, we use the number of lamin reads to model the mean-variance relation for LAD scores. This explained the mean-variance relation better than combined lamin and Dam reads, most likely because Dam signal is more widespread than lamin signal and therefore less informative. We used a linear spline between the time points to call significant changes for the hTERT-RPE time course experiment [52]. Significance was defined as a Benjamini & Hochberg adjusted p-value lower than 0.01. For the cell cycle sorting experiment, we performed pairwise testing on the LAD scores between G1 and G2. The power to detect significant changes was lower because it uses two data points (G1, G2) per replicate instead of five (all the hTERT-RPE time points). To keep the number of differential LADs roughly identical, we decided to change the significance threshold for this experiment (0.05 for K562 and HCT116, 0.001 for Hap1 which has an additional replicate and thus more power to detect significance).

### Active genes definition

RNA-seq reads were downloaded from ENCODE and published reports: HAP-1 ([53], [54], [55], [56]), K562 ([53]), HCT116 ([53]) and RPE ([57]). Reads were processed with fastp 0.12.2 to remove potential adapter sequences and low-quality reads [58] and aligned to GRCh38 v15 and counted in Gencode genes v24 with STAR 2.5.4a [59]. Regularized log_2_ transformed values from DESeq2 1.18.1 were used a measure of gene expression [60]. Active genes were defined as genes with a mean cell score of 7 or higher, which falls between the bimodal distribution of inactive and active genes.

### Replication timing comparison

Processed replication timing data at a 5 kb resolution for hTERT-RPE, K562 and HCT116 cells were downloaded from the 4D Nucleome data repository (https://data.4dnucleome.org). The mean signal of four adjacent bins was used to compare to the 20 kb binned pA-DamID data.

### Telomere and centromere distance

Telomeres were defined as the first and last base of GRCh38. Centromere locations for GRCh38 were downloaded from the UCSC table browser (group: mapping and sequencing; track: centromeres).

### Data availability

Reads and processed data files for the generated pA-DamID and DamID data will be made available via the 4D Nucleome data repository (https://data.4dnucleome.org/).

## Supplementary figure legends

**Figure S1.**
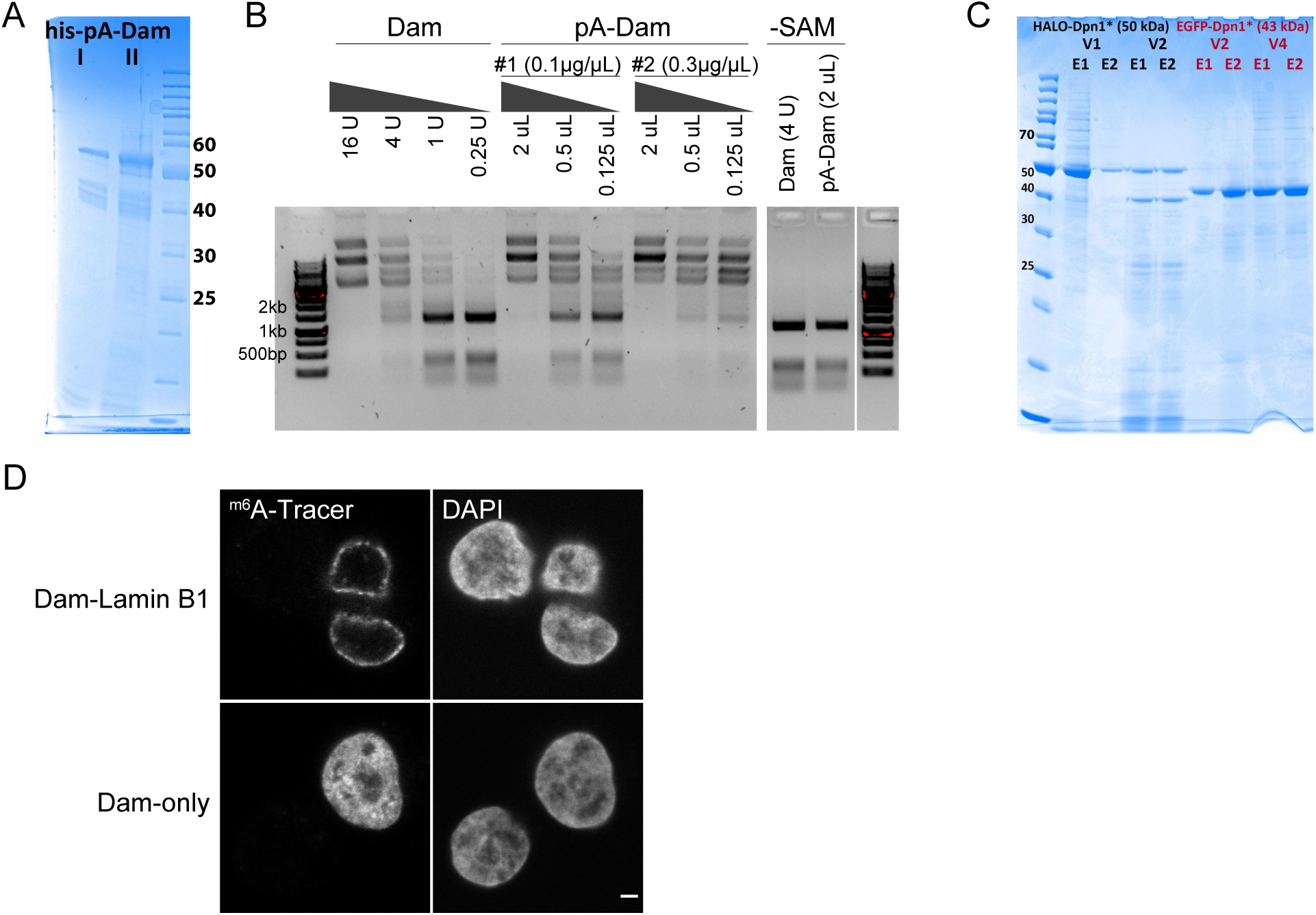
Production and validation of the pA-DamID proteins. (**A**) SDS-PAGE gel of two batches of bacterial purified pA-Dam protein (expected molecular weight: 52 kDa) with concentrations of ∼0.1 and ∼0.3 µg/µL, respectively. (**B**) Agarose gel analysis of Mbo I protection assay. Unmethylated plasmid is ^m6^A methylated by a range of Dam and pA-Dam concentrations. The DNA is subsequently digested by Mbo I, which can cut GATC but not G^m6^ATC sequences. The pA-Dam batches have Dam activities estimated to be ∼8 and ∼32 units (see Methods for definition) per ∼0.1 µg and ∼0.3 µg pA-Dam protein, respectively. No Mbo I protection is observed without the methyl-donor SAM. (**C**) SDS-PAGE gel of ^m6^A-Tracer protein purified from insect cells, with the truncated DpnI fused to GFP and HALO tags. Two elutions (labeled E1/E2) of two independent virus pools (labeled V1/V2 and V2/V4) produced protein of the expected size (50 and 43 kDa for GFP and HALO-tagged protein, respectively) and were pooled. (**D**) Confocal sections of ^m6^A-Tracer signal in HAP-1 cells transduced with lentivirus expression Dam-Lamin B1 (*top panel*) and Dam-only (*bottom panel*). Negative cells are presumably non-transduced cells. The scale bar corresponds to 2 µm.

**Figure S2.**
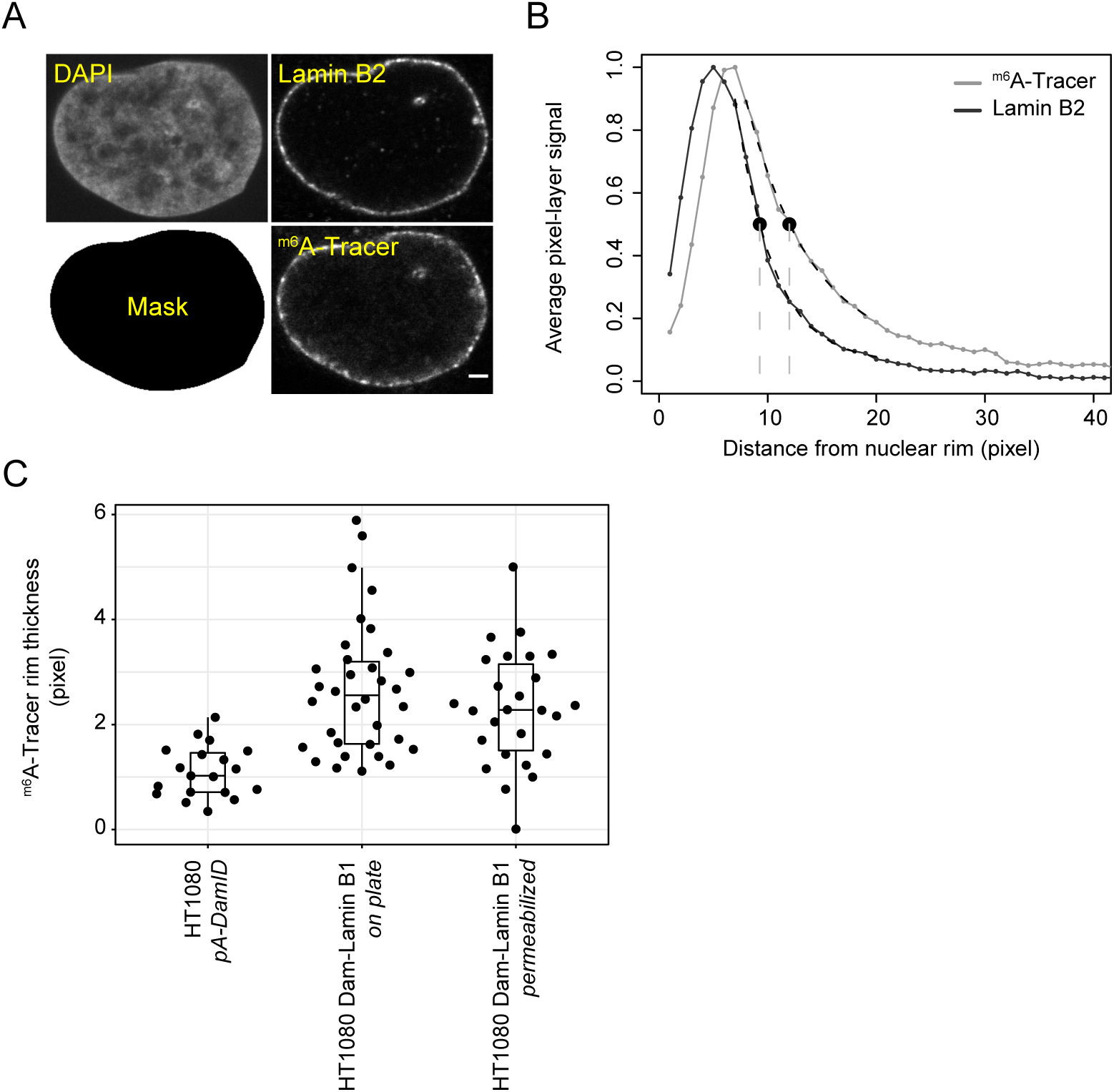
Quantification of ^m6^A-Tracer rim signal for DamID and pA-DamID. (**A**) HT1080 cells expressing inducible Dam-Lamin B1 [8] were treated with Shield1 to induce ^m6^A methylation. Cells were either fixed immediately or processed according to the pA-DamID protocol (as negative controls) and then fixed on poly-L-lysine coated cover slips. Cells were imaged by confocal microscopy for the NL (Lamin B2 antibody) and methylation (^m6^A-Tracer) for both conditions. The scale bar corresponds to 2 µm. (**B**) For every cell, the 50% decay distance from the nuclear periphery was determined for Lamin B2 and ^m6^A-Tracer by fitting exponential decay functions from DAPI-segmented nuclei. The difference in 50% decay distance between Lamin B2 and ^m6^A-Tracer was used as a measure of the thickness of the ^m6^A-Tracer layer. (**C**) Distribution of the ^m6^A-Tracer layer thickness for HT1080 cells expressing Dam-Lamin B1, visualized before and after the pA-DamID protocol. For comparison, a similar analysis was performed with HT1080 cells not expressing Dam-LaminB1, subjected to Lamin B2 pA-DamID.

**Figure S3.**
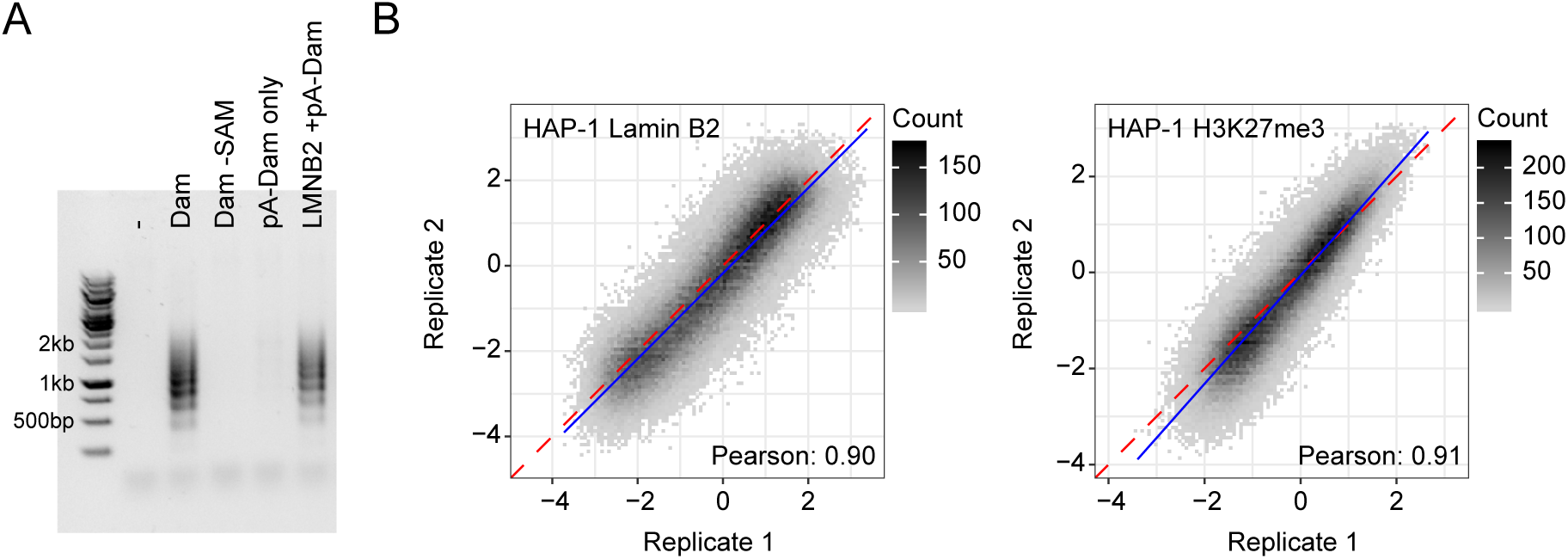
Additional controls for pA-DamID mapping data generation. (**A**) Gel analysis of amplified ^m6^A fragments for a representative experiment. DNA isolated from pA-DamID processed cells was digested with DpnI and fragments were ligated to PCR adapters. After 15 PCR cycles, a sample is loaded on gel to visualize amplification. A negative control and no-SAM controls samples do not yield any amplification product, as expected. pA-Dam without primary antibody gives a weak background amplification, but much less than targeted pA-Dam. (**B**) Genome-wide correlations of pA-DamID data (bin size 20 kb) between two biological replicates in HAP-1 cells for Lamin B2 (*left panel*) and H3K27me3 (*right panel*).

**Figure S4.**
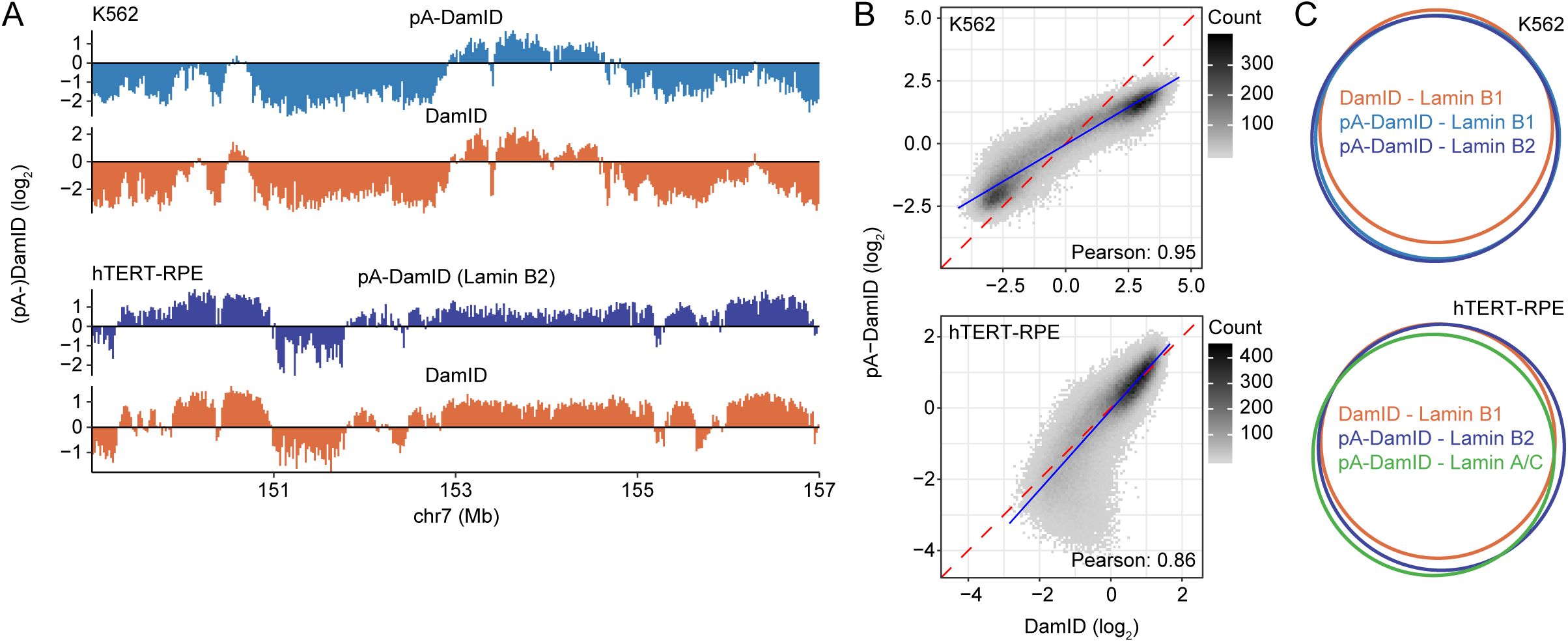
Comparison of NL interactions determined by pA-DamID and DamID for K562 and hTERT-RPE cells. (**A-C**) Similar plots to Fig. 2A-C for K562 and hTERT-RPE cells. Because we did not generate Lamin B1 data for hTERT-RPE cells, Lamin B2 data is shown instead.

**Figure S5.**
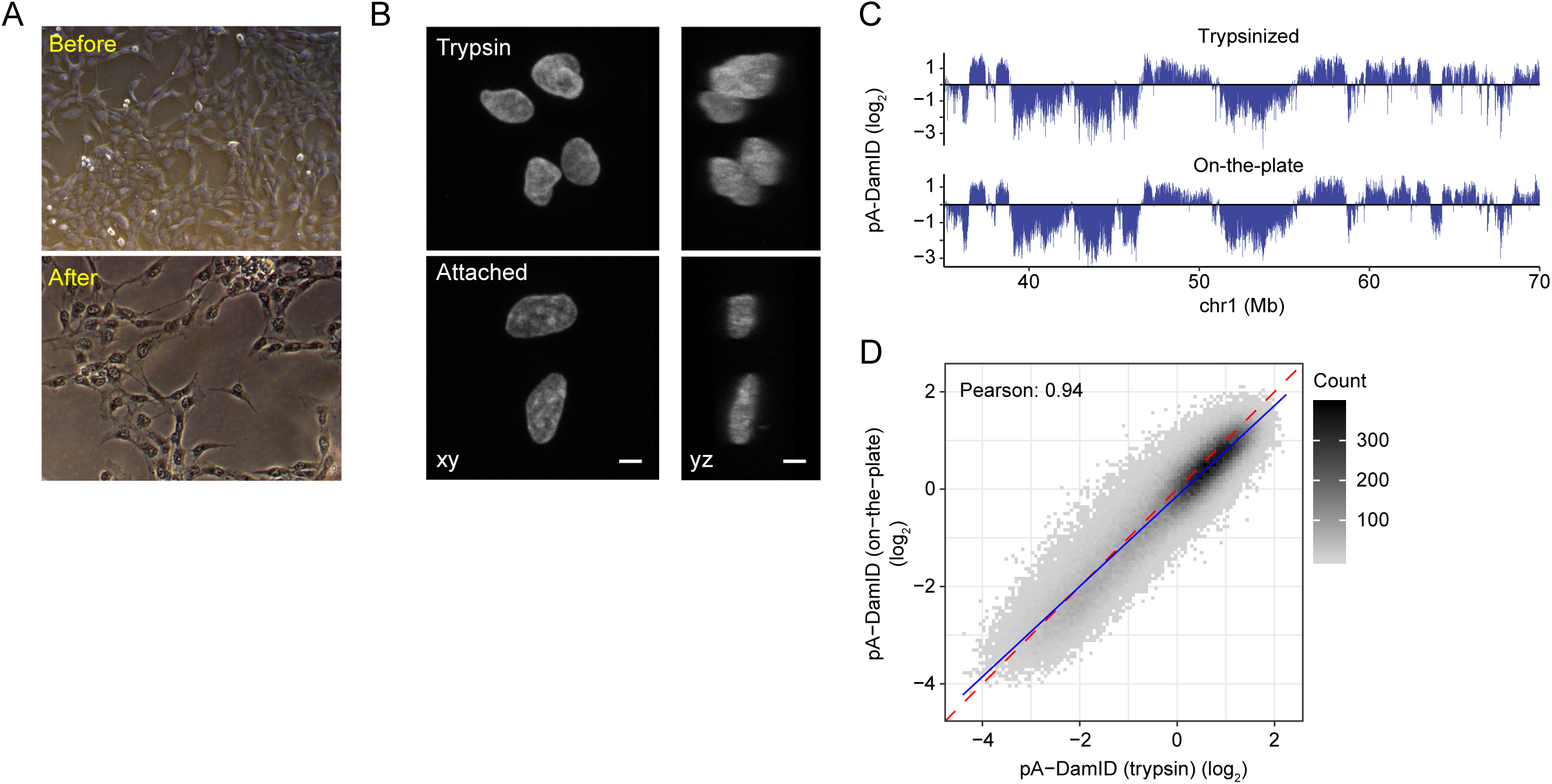
The effect of nuclear shape on NL interactions. (**A**) Light microscopy of hTERT-RPE cells attached to the culture dish before and after pA-DamID protocol. (**B**) Confocal sections visualizing the 3D shapes (xy plane and interpolated yz plane) of DAPI-stained nuclei of trypsinized cells and of cells attached to the culture dish. Trypsinized cells are rounder than attached cells. Scale bar corresponds to 5 µm. (**C**) Representative locus comparing pA-DamID data tracks obtained with trypsinized cells and attached cells. Shown are the averaged signals of two replicates, bin size 20kb. (**D**) Genome-wide correlation of pA-DamID data (bin size 20 kb) between trypsinized and adherent (on-the-plate) cells.

**Figure S6.**
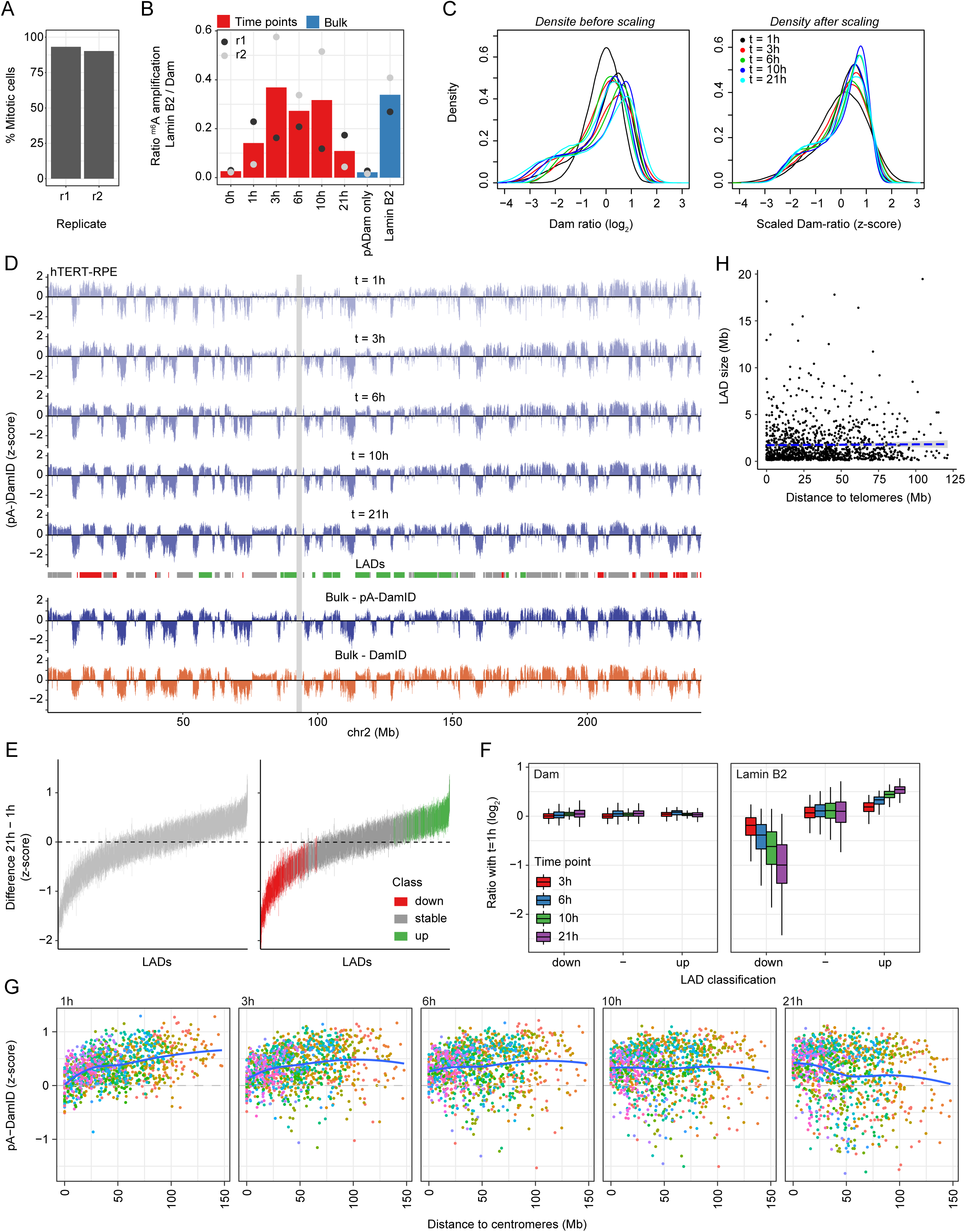
Analysis of NL interactions at various times after mitosis. (**A**) Synchronized cells were fixed on poly-L-lysine coated cover slips and stained with DAPI. Efficiency was estimated by manual counting of metaphase and anaphase appearance in confocal microscopy images. 207 and 82 cells were assessed in the two replicates, respectively. (**B**) Ratio of gel intensities from amplified ^m6^A fragments between Lamin B2 pA-DamID and Dam control, normalized for input DNA. For the second replicate, input DNA was very low for the timepoints and could not be determined accurately. These samples thus show a large variation in amplification ratio compared to the first replicate. The 0h time point consistently shows low amplification ratios, similar to bulk samples without a primary antibody. This indicates that genome – NL interactions at this time point are very weak. (**C**) Data distribution before and after conversion to z-scores. The distributions are not affected by this transformation. (**D**) Similar plot as Fig. 4B, showing a complete chromosome (chr2). LADs are colored by their differential class (down:red, stable:grey, up:green, see Fig. S6E). The centromere is shown as grey box. (**E**) Overview of the classification of dynamic LADs. LADs were defined for every time point by selecting LADs that are called in both replicates using hidden Markov modeling, after which a union across time points was used as consensus LAD definition. For every LAD, per-bin z-score differences are calculated between 1h and 21h, and a line is drawn between the 25^th^ and 75^th^ percentiles of these values. This shows that numerous LADs change as complete units between 1h and 21h. The right panel shows the same figure, but is colored by LAD classification into stable and dynamic using limma-voom differential analysis (Methods). (**F**) Read counts for Dam and Lamin B2 (normalized for overall sequencing read depth) were calculated for stable and dynamic LADs. Plotted are log_2_-ratios relative to the 1h time point. This shows that Dam reads do not change between time points. (**G**) Similar figure as Fig. 3I, showing distance to centromeres instead. (**H**) LAD size is not correlated with distance to telomeres (Pearson correlation coefficient: 0.01).

**Figure S7.**
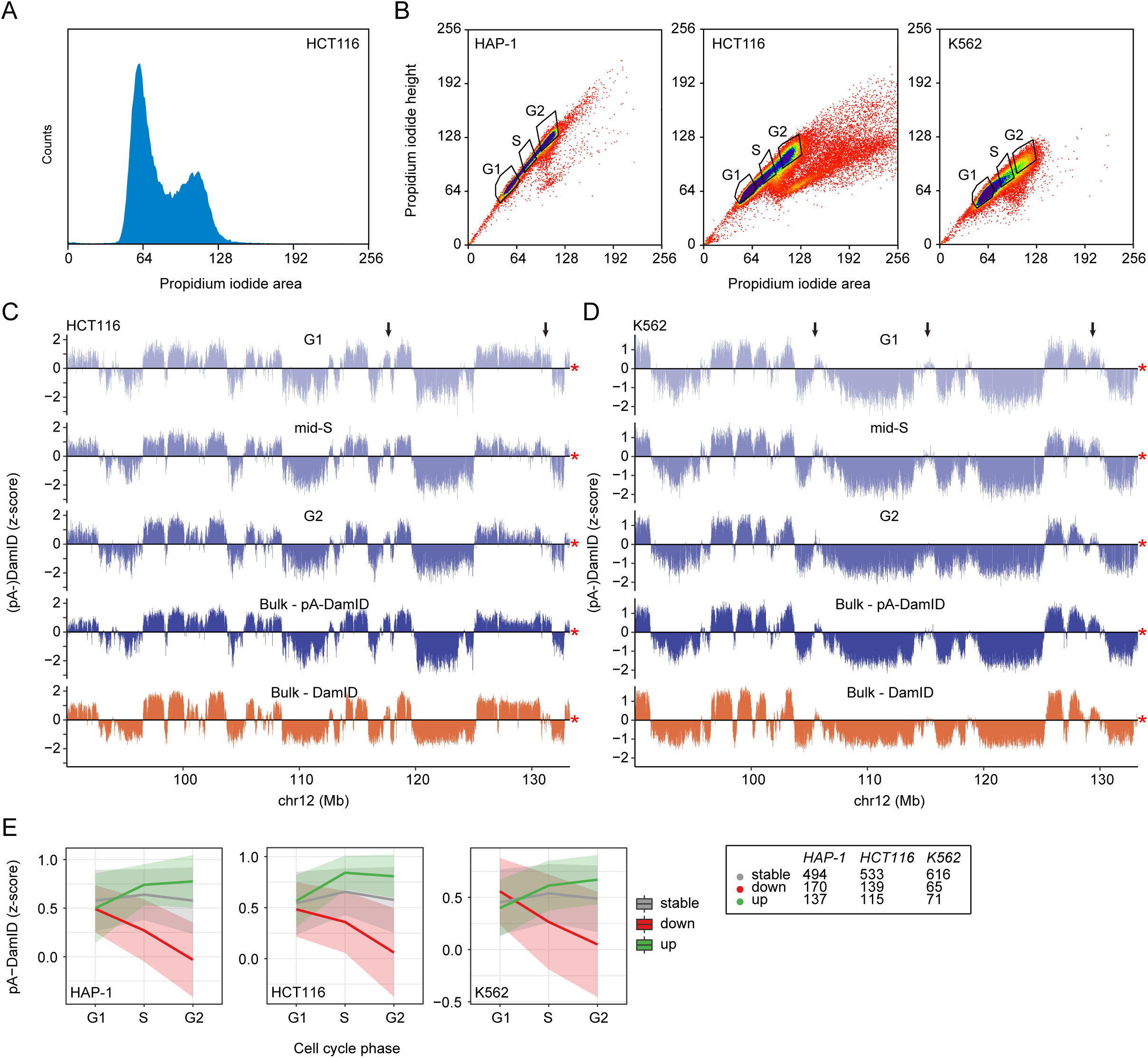
LAD dynamics in flow sorted HCT116 and K562 cells. **(A)** Density plot of propidium iodide (PI) signals of HCT116 cells after application of the pA-DamID protocol and PI staining. (**B**) Sorting strategy of pA-DamID processed cells stained PI. Single cells were selected based on PI height and area and sorted into three pools: G1, mid-S and G2. G1 and G2 gates were set around the two PI peaks (see also A). The mid-S gate was set to covering intermediate PI signals, clearly distinct from the G1 and G2 peaks. For every condition, 2 or 3 wells were filled with 1000 cells and processed to generate one sequencing library. (**C-D**) Same locus as Fig. 4A for HCT116 (**C**) and K562 (**D**) cells. Arrows highlight dynamic regions. (**E**) Similar plot as Fig. 3C, with cell cycle stages on the x-axis. The numbers of stable and dynamic LADs are shown for the three cell types.

**Figure S8.**
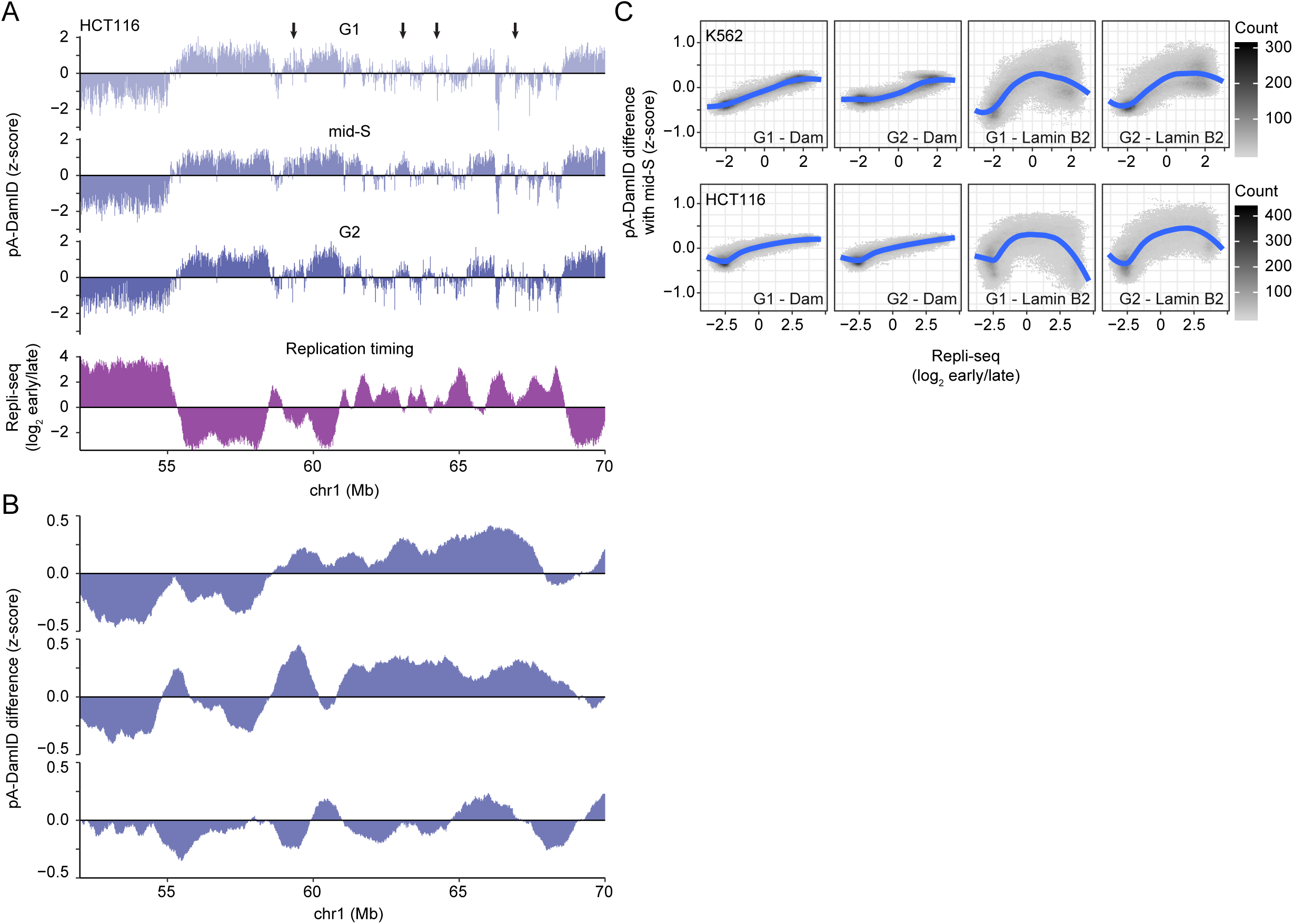
Transiently increased Lamin B2 interactions during mid-S are not due to increased Dam reads. (**A-B**) Similar figures as Fig. 5A-B for a locus in HCT116 cells. Arrows highlight regions with increased mid-S signal compared to G1 and G2. (**C**) Similar plot as Fig. 5C, showing the difference in read-normalized counts for Dam and Lamin B2 samples. This plot shows a gradual increase in S-phase enrichment for Dam reads, reflecting DNA duplication of early replicating DNA. In contrast, Lamin B2 reads show an increase in S-phase enrichment for Repli-seq scores around 0, reflecting mid-S chromatin.

**Figure S9.**
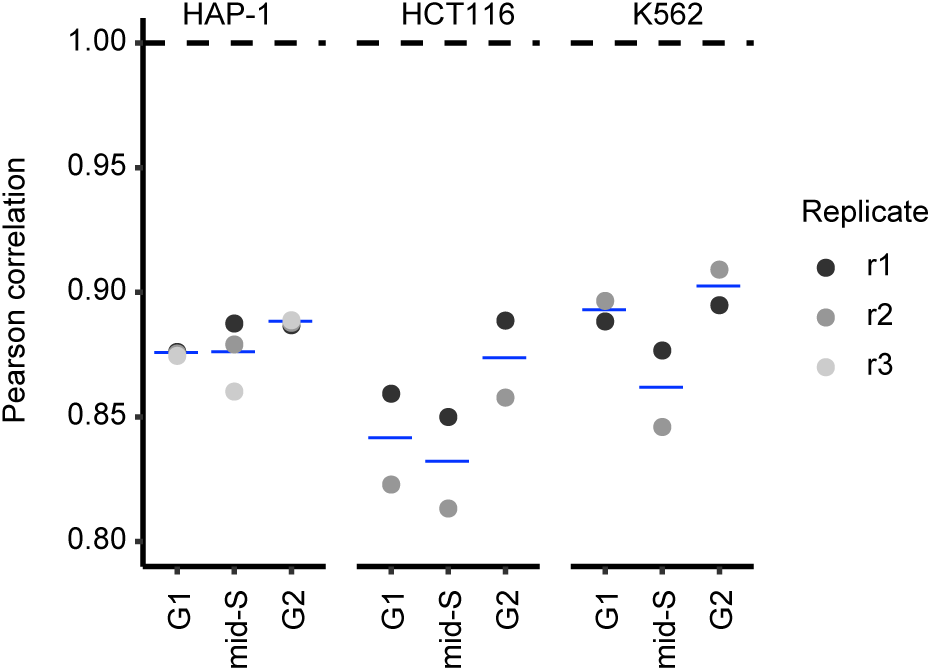
Correlation of cell cycle sorted pA-DamID with DamID. (**A**) Pearson correlation with DamID signal values are shown for every cell cycle stage. Correlations are consistently highest for G2 cells. This is most apparent for HCT116 cells, that showed the biggest difference between pA-DamID and DamID data.

## Supplementary information

**Supplementary Information S1.**

Detailed protocol of pA-DamID.

