## Supplementary Information S1 for "Cell cycle dynamics of lamina associated DNA"

### pA-DamID (TS)

Owner: Tom van Schaik | Created at: November 29, 2018

#### Protocol description

https://data.4dnucleome.org/experiment-types/pa-damid/#overview

Experiment category: Sequencing
Assay classification: Linear DNA enrichment
Assay purpose: Proximity to Cellular Component
Raw files: Reads (fastq) provided by lab

Short description:

The proteinA-DamID (pA-DamID) method is a hybrid of the CUT&RUN {Skene, 2017, 28079019} and DamID {Vogel, 2007, 17545983} technologies. Cells are permeabilized with digitonin and incubated with an antibody against a nuclear protein of interest, followed by binding with pA-Dam. Subsequent activation of the tethered Dam by its methyldonor S-adenosyl-methionine (SAM) results in *in situ* G$^{m6}$ATC methylation of DNA sequences that are in molecular proximity to the protein of interest. The pattern of deposited $^{m6}$A can be mapped genome-wide as in conventional DamID.

The labeled DNA can also be visualized using the$^{m6}$A-Tracer {Kind, 2013, 23523135}, providing a powerful quality control and possibly new biological insights. In addition, labeled cells can be sorted to study (rare) subpopulations of interest.

#### pA-DamID protocol

pA-DamID protocol to probe snapshot protein-DNA interactions using the general DamID procedure. This protocol covers the parts from cell culture to SAM incubation. After this, three options are possible (or a combination of them) :

1. Bulk genomic DNA isolation, followed by the bulk DamID protocol.
2. m6A-Tracer binding, followed by fixation on polylysine slips.
3. Single-cell sorting, followed by the single-cell DamID protocol.

This labguru protocol is based on CUT&RUN and initial rounds of modifications, see "ts180820_pADamID".

#### Reagents & preparation

Reagents are prepared before starting the procedure.

**5% digitonin** - prepared fresh:

- Solubilize digitonin in preheated (~95C) MQ: 10mg / 200uL
  - Note: "preheat" your pipet tip by going up and down a few times

**Wash buffer** - prepared fresh:

- 1 mL 1M HEPES-KOH pH=7.5 (stored at 4C)
- 1.5 mL 5M NaCl (29.2 g / 100 mL $H_2O$)
- 25 uL 1M spermidine (1M = 0.145 g/mL)
  - Note: you can a store an aliquoted stock solution at -20C, for ~3 months max.
- 1 Roche Complete tablet -EDTA

- H2O up to 50 mL

**Dig-Wash buffer** - prepared fresh:

- 200-1000 uL 5% digitonin to 50 mL wash buffer (0.02% - 0.1%)
    - Note: too much digitonin will also disrupt your nuclear membrane and compromise your nuclear integrity
    - Note: for most cell types, 0.02% is plenty

**Activation buffer** - prepared fresh:

- Dig-Wash buffer, supplemented with 1/400 SAM (32 mM stock, 80 uM working solution)
    - Note: no need to make this one in advance

While working, buffers are kept on ice.

A centrifuge is pre-cooled to 4C.

### 1. Nuclear isolation

1. Prepare cells. (~1-2M / condition is a very safe starting amount.)
2. Wash cells with PBS or Wash buffer, spin and remove supernatant, 3 minutes at 500g.
3. Wash cells with Dig-Wash buffer, spin and remove supernatant.
    - Note: at this point, split your bulk pool in individual samples.

Import controls to add:

1. Completely negative control.
2. Negative control, adding Dam enzyme during the activation.
3. Most important: no primary antibody control, probing DNA accessibility + amplification bias.

### 2. Bind primary antibody

1. Gently resolve pellet in 200 uL of Dig-Wash (per sample) containing your antibody.
    - Either premix your antibody with Dig-Wash, or add the antibody afterwards followed by careful mixing.
    - Antibody concentration should be optimized. 1:100 is a good starting concentration for strong signal, while 1:500 is a good concentration for a higher dynamic range.
2. Place on the rotator at 4C for ~2h.
3. Spin and remove supernatant.
4. Wash once with 0.5mL of Dig-Wash buffer.
    - Be careful not to lose nuclei on the side of the tube, especially when working with few nuclei. We usually spin for 2 minutes, followed by half a twist and another minute to prevent nuclei lining on the side of the tube.

### - Optional: bind secondary antibody

1. Gently resolve pellet in 200 uL of Dig-Wash (per sample) containing a bridging antibody (i.e. 1:100 rabbit anti-mouse bridging antibody).
2. Place on the rotator at 4C for ~1h.
3. Spin and remove supernatant.
4. Wash once with 0.5mL of Dig-Wash buffer.

### 3. Bind pA-Dam

1. Gently resolve pellet in 200 uL of Dig-Wash (per sample) containing ~ 1:100 pA-Dam.
    1. Note: pA-Dam concentration can be varied and depends on the protein batch.
    2. **Control:** add pA-Dam to an otherwise negative sample.
2. Place on the rotator at 4C for ~1h.
3. Spin and remove supernatant.
4. Wash with 0.5mL of Dig-Wash buffer. In total 2 times.

### 4. Activation

1. Gently resolve in 100 uL of the Activation Buffer (per sample).
2. Incubate at 37C for 30'.
    1. **Control:** add Dam enzyme to an otherwise negative sample as Dam-only control.
3. Spin and remove supernatant.
4. Wash once with Dig-wash buffer.

### What's next

At this point, cells are permeabilized and labeled with $^{m6}$A modifications near the protein of interest. Three options are possible at this point:

1. **Genome-wide mapping of $^{m6}$A modifications.** To generate genome-wide mapping data, follow the library preparation for conventional DamID (https://data.4dnucleome.org/experiment-types/damid-seq/). However, the DpnII digestion must be omitted (see note below), and 4 uL of DpnI adapter ligation product can be used in 40 uL total mePCR reaction. Amplification requires between 15-19 PCR cycles for most samples.

   Note: It seems that pA-DamID is not able to methylate on nucleosomal DNA. This is not an issue for DamID, where Dam is expressed in live cells for prolonged periods and nucleosomes can remodel to allow Dam to access all DNA. To prevent high losses due to unmethylated GATC sequences between methylated GATC sequences, omit the DpnII digestion for pA-DamID samples.

2. **Visualization of $^{m6}$A modifications.** To visualize the position of protein-DNA interactions (indicated by $^{m6}$A modifications), follow the protocol "pA-DamID microscopy slides".

3. **Sorting of processed cells.** pA-DamID processed cells can be sorting for specific subpopulations. We have performed propidium iodide staining (see protocol "pA-DamID cell sorting") to gather G1, mid-S and G2/M subpopulations.

### Reagents

- Digitonin: Millipore, #300410-250MG
- HEPES-KOH: Sigma, #H4034-100G
- NaCl: Sigma, #S5886-10KG
- Spermidine: Sigma, #S0266-5G
- cOmplete™ Protease Inhibitor Cocktail EDTA-free: Roche, #11873580001
- Rabbit anti-mouse bridging antibody: Abcam, #ab6709
- Rabbit anti-goat bridging antibody: Abcam, #ab6697
- S-adenosylmethionine (SAM): New England BioLabs, #B9003S
- Dam enzyme: New England BioLabs, #M0222L
- pA-Dam: this manuscript

### pA-DamID microscopy slides (TS)

Owner: Tom van Schaik | Created at: November 29, 2018

#### Protocol description

pA-DamID results in intact, permeabilized cells with $^{m6}$A deposited near the antibody of interest. To confirm that the m6A modification is deposited at the expected site or to study subpopulations with varying levels / patterns of m6A modification, it is valuable to visualize the pattern of $^{m6}$A modifications. For example, when performing pA-DamID for the nuclear lamina, the peripheral enrichment of $^{m6}$A modifications correlates with the quality of the data set. This protocol describes the staining of $^{m6}$A modifications using isolated $^{m6}$A-Tracer protein {Kind, 2013, Cell}.

#### pA-DamID microscopy slides

After pA-DamID, $^{m6}$A modifications can be visualized inside the intact nuclei using the $^{m6}$A-Tracer protein (coupled to GFP and HALO). This can be done in two ways:

1. Bind the nuclei to poly-L-lysine-coated coverslips, fixate and bind with the $^{m6}$A-Tracer.
2. Bind the $^{m6}$A-Tracer first, and then bind on poly-L-lysine-coated coverslips and fixate.

This procedure is written for possibility #2, but can easily be adopted for #1. Furthermore, this protocol uses buffers from the pA-DamID protocol although these can be replaced with more generic alternatives.

#### 1. Bind m6A-Tracer

Starting material: pellet of washed nuclei with $^{m6}$A modifications.

1. Gently resolve pellet in 100 uL of cold Dig-Wash (per sample) (or PBS) containing ~1:1000 $^{m6}$A-Tracer and possibly a secondary antibody.
2. Place on the rotator at 4C for 30 - 60 minutes.
3. Wash with Dig-Wash buffer.

#### 2. Bind nuclei to poly-L-lysine-coated coverslips

1. Resolve nuclei in ~300 uL Dig-Wash and place on 0.1%-coated poly-L-lysine-coated cover slips.
2. Incubate for 30 minutes on ice.

#### 3. Fix and mount

1. Wash with cold PBS, very gently. Bring to room temperature.
2. Fixate with 2% room temperature formaldehyde/PBS for 10 minutes.
3. Wash with PBS. Wash with Demi water.
4. Mount with Vectashield + DAPI and dry.
5. Seal with nail polish.

#### Reagents

- DigWash: Digitonin Wash buffer from pA-DamID protocol

- Poly-L-lysine: Sigma-Aldrich, #P8920-100ML
- Vectashield+DAPI: Vector Laboratories, #H-1200
- $^{m6}$A-Tracer: {Kind, 2013, Cell}; isolated protein in this manuscript

### pA-DamID cell sorting (TS)

Owner: Tom van Schaik | Created at: December 17, 2019

#### Protocol description

DamID library preparation requires little input material and has already been implemented for single-cell sequencing {Kind, 2015, Cell}. As pA-DamID is a technical modification of DamID, this also applies to pA-DamID. pA-DamID results in intact, permeabilized cells with $^{m6}$A modifications that can easily be sorted using flow cytometry. Using this modified single-cell protocol, $^{m6}$A modifications can be map genome-wide for (rare) subpopulations.

This modified single-cell protocol is specific for propidium iodide staining of pA-DamID processed cells and sorting based on DNA content, resulting in 1000 cells per well.

#### General guidelines

Low amounts of starting material are more susceptible to sample contamination. To prevent this, work in a clean environment. For us, this means:

- Work in the PCR cabinet until first methyl PCR
- PCR cabinet cleaned with ZAP and UV
- Use the special single cell pipet set in the PCR cabinet up until first methyl PCR
- Use Clean epjes from separate bag, and filter tips
- Keep separate stock solutions for all the single cell work
- Do not move your hands over the PCR plate
- Number and date you plates

#### 1. FACS sorting

**Preparation for the sorting**

1. Prepare 96 wells PCR plates with 3ul **1.33x** lysis buffer + ProtK in each well. Use repeating pipet to dispense the lysisbuffer.
    - Mastermix per plate:
        - 300 ul 1.33x Single Cell lysisbuffer
        - 17.7 ul protK (Roche #03115887001 pH7.5 18 +/- 4 mg/ml)
    - Quick spin plate after, cover with Microseal A film (#MSA5001 Biorad) put them on ice
2. Resuspend cell pellet in 1 mL low serum containing medium (PBS + 2% FBS + 1:500 propidium iodide), put in FACS tube and place on ice.
3. Together with the plates move down to the FACS.

**Cell cycle sort modification: This is a modified single-cell DamID protocol {Kind, 2015, Cell}. Note that for these experiment we sorted 1000 cells into a single well. For these numbers, volumes start to become relevant as 1000 cells is roughly 1uL. This requires 1.33x lysis buffer because 3uL + 1uL sorted cells = ~4uL, at which point the 1.33x should be back at 1x.**

**Sorting strategy**

1. For haploid cells, gate on small cells in the population.
2. Set gates based on propidium iodide signal.
3. Sort cells according to the following template:

**Cell cycle sort - plate format**

| | 1 | 2 | 3 | 4 | 5 | 6 | 7 | 8 | 9 | 10 | 11 | 12 |
|---|---|---|---|---|---|---|---|---|---|---|---|---|
| A | Dam - G1 -1000 | Dam - S -1000 | Dam - G2 -1000 | LMNB2 - G1 -1000 | LMNB2 - S -1000 | LMNB2 - G2 -1000 | | | | | | |
| B | | | | | | | | | | | | |
| C | | | | | | | | | | | | |
| D | Dam - G1 -1000 | No cells | | LMNB2 - G1 -1000 | No cells | | | | | | | |
| E | Dam - G1 -1000 | | | LMNB2 - G1 -1000 | | | | | | | | |
| F | | | | | | | | | | | | |
| G | | | | | | | | | | | | |
| H | | | | | | | | | | | | |

Note that wells D1/4 and E1/4 can be used a control wells (-DpnI / -Ligase) and the "no cells" as H2O controls.

#### 2. Lysis

1. After sort, quick spin the plates, and incubate 4hr at $53^{0}$C in a PCR machine.
2. 10' 80 $^{0}$C heat inactivate (PCR machine)
3. Quick spin
4. Before continuing, carefully remove top (melts a bit unto plate at 80 degrees)

You can either store the plates at -20 now, or before the heat inactivation or immediately proceed with the rest of the protocol. Upon thawing plates from the -20 always check if there is still sufficient liquid in all wells after quick spin. If not adjust with nuclease free water until approximately 4 uL.

#### 3. DpnI digestion

Per reaction

- 0.7ul phos all buf
- 0.1ul DpnI
- 0.07ul 10% Tween-20
- 6.13ul MQ

Make mastermix for 1 plate (100x)

- 70ul phos all buffer
- 10ul Dpn1
- 7ul 10% Tween-20

- 613ul MQ

1. Add 7ul/well digestion mix using repeating pipet
2. Incubate 4hr 37$^0$C, followed by a quick spin
3. Heat inactivate 20’ 80$^0$C in a PCR machine, followed by a quick spin

#### 4. DpnI adapter ligation

Per reaction

- 2ul T4 ligase buffer
- 0.5ul ligase (5U/ul)
- 0.1ul annealed ds adaptor (50uM)
- 7.4ul MQ

Make mastermix for 1 plate (100x)

- 200ul 10x ligase buffer
- 50ul ligase (5U/ul)
- 10ul Dpn1 adaptor
- 740ul MQ

1. Add 10ul/well ligation mixture using repeating pipet
2. Incubate 16hr at 16 degrees
3. Heat inactivate 10’ 65, followed by a quick spin

#### 5. Methyl PCR

Per reaction

- 25ul MyTaq
- 1.0 ul primer Adr-PCR (Jop 528; 50uM)
- 4.0 ul MQ

make mastermix, for 1 plate (110x)

- 2750 ul MyTaq
- 110 ul Adr-PCR (528)
- 440 ul MQ

1. Using repeating pipet 30ul/well (4ul tip, step 7 1/2), pipet in with some force to mix well quick spin
2. PCR according to protocol below

| **1 cycle** | **1 cycle** | **4 cycli** | **~17 cycli** | **1 cycle** |
|---|---|---|---|---|
| 10’ 72˚C | 1’ 94˚C | 1’ 94˚C | 1’ 94˚C | 10’ 72˚C |
| | 5’ 58˚C | 1’ 58˚C | 1’ 58˚C | forever 4˚C |
| | 15' 72˚C | 10' 72˚C | 2' 72˚C | |

3. Final set of cycli ~17x (to be optimized)

4. From now on no need for PCR cabinet anymore

### 6. Load on gel

1. After PCR, pipet 4ul of PCR onto maxi 1% gel
2. Next to 4ul marker, run 30’ at V 160 V, 400mA

### 7. PCR products clean up using CleanPCR beads

Technical replicates are pooled together and PCR products cleaned using 1.8x CleanPCR beads.

**Steps**

| | |
|---|---|
| 1 | Prewarm the CleanPCR beads 10’ at RT |
| 2 | Pipet beads slowly, and keep resuspending before you take out |
| 3 | Add 1.8x sample-volume of the beads = 90µl |
| 4 | mix by pipetting 10x |
| 5 | Let bind for 5’ at RT |
| 6 | Place tubes in the tube magnetic rack and allow the beads to move to the magnetic side (approx. 3’) |
| 7 | Remove supernatant |
| 8 | Wash with 500µl icecold 80%EtOH by pipetting not directly on the beads but on the side of the tubes whilst they remain in the rack, |
| 9 | Incubate 30” |
| 10 | Aspirate the EtOH in 2 goes and repeat wash once (2x in total) |
| 11 | After final aspiration let the beads dry for a bit (~5 minutes) |
| 12 | Elute the material from the beads with 25 uL nuclease free $H_2O$ (RT) |

| | |
|---|---|
| 13 | Mix by pipetting 10x |
| 14 | Incubate 3’ RT |
| 15 | Place in magnetic rack and let beads separate for 3’ |
| 16 | pipet solution over in clean tube, be careful not to take along beads |

#### 8. Blunt ending 3’ overhang with End repair kit

Make mastermix of

5µl 10x end repair buffer

5µl dNTP

5µl ATP

9µl nuclease free $H_2O$

1µl end repair enzyme mix

***everything from end repair kit (except $H_2O$)***

add 25µl to each sample

incubate 45’ RT on bench

(no heat inactivation)

Instead, bead purification as step #7, elute in 26 µl preheated (50C) nuclease free H2O.

#### 9. Addition of 3’-A overhang

Prepare fresh ice cold 80% EtOH for bead purification later on

Correct elution volume to 25 µl

Make mastermix of

5µl NEB buffer 2

0.1µl 100mM dATP (! Not regular ATP)

19.4µl nuclease free $H_2O$

0.5µl Klenow fragment NEB (3’ 5’ exo-)50U/µl

add 25µl mastermix with Gilson 1 by 1 to the samples and mix by pipetting up and down, immediately place on block heater at 37ºC (to avoid strange reactions of Klenow at different temp) incubate 30’ 37ºC

heat inactivate 20’ 75ºC

#### 10. CLeanPCR bead cleanup

Bead purification as step #7.

Nanodrop and correct for the difference in concentration between the different samples. Adjust all samples to approximately 40ng/µl if possible.

#### 11. Ligation of illumina Y-adaptor

Add to 220 to 250 ng of PCR products (in 6.5 µl ), 3.5 µl of ligation master mix.

As a control take along >1 no ligase sample

Ligation mastermix

- 0.5 µl ds Y-Adaptor (50 mM)
- 1 µl 10x ligation buffer
- 0.5 µl T4DNA ligase (5U/ml)
- 1.5 µl $H_2O$

final volume 10ml

Incubate 16h at 16ºC

inactivate 10’ 65˚C

#### 12. CleanPCR bead cleanup

Add 40µl nuc free $H_2O$ and transfer to an tube.

purify using 1.8x volume beads and magnet as before (step 10), elute with 20µl nuclease free $H_2O$.

#### 13. Index PCR (add the indices)

Pipet in PCR strip

- 8 µl DNA with Y adaptor (8µl roughly 100ng)
- 0.5 µl Illumina index primer (10mM) a different one in every reaction

Index PCR master mix

- 1 µl $H_2O$
- 0.5 µl P5-Illumina-2 PCR (10mM)
- 10 µl 2xMyTaq mix

Total volume 20 µl

Program:

| 1 cycle | 9-12 cycles(**) | 1 cycle | forever |
|---|---|---|---|
| 1’ 94 ºC | 30” 94ºC | 2’ 72ºC | 4ºC |
| | 30” 58ºC | | |
| | 30” 72ºC | | |

(**) (number of cycle to optimize, enough to get a visible smear on gel, but as few as possible as too much product may affect sequencing efficiency)

#### 14. Check on gel

Load 4 µl of PCR and 1.8µl of input (same amount taking into account dilution in PCR mix). The bulk of the smear should be below 1.5 kb.

#### 15. Pool the samples

Based upon smear intensity on gel. Determine the final volume.

#### 16. Bead purify

Now use 1.6x bead volume vs sample

elute in 20 µl nuclease free $H_2O$ and nanodrop.

Expect to have between 400-700 ng/µl.

Run sample on gel to make sure all primers are removed, if not do Bioline Isolate II PCR and Gel column purification or cut the appropriate size fraction from gel and purify if the smear fragments are too high in size. (Ideally they should be between 300-1500bp)

#### Reagents


**Primer sequences (not phosphorylated) from IDT:**

p528 DpnI PCR NNNNGTGGTCGCGGCCGAGGATC

p530 DpnI adapter_bottom TCCTCGGCCGCG
p532 Adaptor_top CTAATACGACTCACTATAGGGCAGCGTGGTCGCGGCCGAGGA

Y-adaptor top ACACTCTTTCCCTACACGACGCTCTTCCGATCT

Y-adaptor bottom **P-**GATCGGAAGAGCACACGTCT
**(5' phosphorylated)**

Ill2-p5 primer AAT GAT ACG GCG ACC ACC GAG ATC TAC ACT CTT TCC CTA CAC GAC GCT CTT CCG ATC T

Keep separate stocks for single cell use only!
Store at RT

**Lysis buffer, final concentrations (1x)**:
10mM Tris.acetate pH7.5 (Sigma 93337-25mg)
10mM Mg.acetate (Sigma 63052-100ml, 1M)
50mM K.acetate (Sigma 95843-100ml-F, 5M)
0.67% Tween 20 (Sigma P1379-25ml)
0.67% Igepal (Sigma I8896-50ml)
0.67mg/ml ProtK (Roche #03115887001 pH7.5; 18 +/- 4 mg/ml)

Prepare by mixing together in a Falcon tube:
1.0ml 0.5M Tris Acetate pH 7.5
0.5ml 1M Mg Acetate
0.5ml 5M K Acetate
335ul Ipegal
335ul Tween20
47.23ml Nuclease free water

add ProtK fresh just prior to use

**10* One-Phor-all-buffer-plus, final concentrations:**
100mM Tris.Acetate, pH7.5
100mM Mg Acetate
500mM K Acetate

Prepare 50ml by mixing together in a Falcon tube:
10ml Tris Acetate (0.5M pH7.5)
5ml Mg Acetate (1M)
5ml K Acetate (5M)
30ml nuclease free water

The 0.5M Tris acetate you prepare by dissolving 3.6238g in 30ml of nuclease free water, then add 1.65ml 1N NaOH (1.599 gram in 40ml) and check the pH to be 7.5

**Adaptor annealing buffer, final concentrations:**
10mM Tris (pH 7.5-8.0)
50mM NaCl
1mM EDTA

mix together in a 50ml Falcon
0.5ml 1M Tris pH7.5-8.0
0.5ml 5M NaCl

0.1ml 500mM EDTA

Perform adaptor annealing according to the following protocol
http://www.sigmaaldrich.com/technical-documents/protocols/biology/annealing-oligos.html

**-20$^0$C**
ProtK Roche #03115887001 pH7.5 18 +/- 4 mg/ml
DpnI NEB #R0176L
T4 DNA Ligase Roche #10799009001
dNTP set Roche #03622614001
Klenow 3′ → 5′ exo– (New Engeland Biolabs #M0212M)
MyTaq Bioline #BIO25043
End-it Repair kit Epicentre #ER81050
